# Hicberg: Reconstruction of contact signals from repeated elements

**DOI:** 10.1101/2025.06.20.660295

**Authors:** Sébastien Gradit, Samuel Ortion, Pauline Larrous, Maëlys Delouis, Romain Koszul, Axel Cournac

## Abstract

In the course of their evolution, genomes can acquire various repeated elements, such as transposons, ribosomal DNA, duplicated genes or tandem repeats. These types of sequences cannot be processed directly by current high-throughput sequencing pipelines, as they generate short reads that cannot be unambiguously localized on reference genomes. We propose an algorithm called *Hicberg* that uses statistical inference with the computation of probability distributions to precisely reassign the positions of reads from repeated sequences in different paired omics data, such as Hi-C data. We show that Hicberg can generate new insights into the impact of repeated elements on the spatial organisation of genomes.

**Significance Statement:** The genomes of microorganisms can contain various types of repeated sequences: duplicated genes, low-complexity sequences and transposons. The question of their potential impact on the spatial organization of genomes is now wide open. We propose Hicberg, an algorithm capable of reconstructing contact signals from repeated elements.

It computes statistical trends on the unambiguous part of the genome and then, by statistical inference, reassigns the position of multi-mapping reads.

The complete chromosome contact maps thus reveal new observations on the impact of repeated elements on chromosome architecture. In particular, they suggest the involvement of certain retrotransposons in the positioning of cohesins, the molecular motors behind chromosome loops.

**Classification:** Biophysics and Computational Biology section

## Introduction

The spatial organization of chromosomes or whole genomes involves different levels of structure, such as chromosomal domains or loops, which can be linked to the cell’s biological functions. In parallel with super-resolution microscopy technologies, chromosome conformation capture (3C) approaches coupled with high-throughput sequencing (Hi-C) have been developed to explore the spatial organization of a genome at different scales (1,2). These technologies are based on the capture of DNA fragments that are in physical contact by fixation with a crosslinking agent combined with high-throughput sequencing to obtain the contact frequencies of different loci within a genome through the formation of chimeric DNA molecules (2). Such methods allow the quantification of contacts between different loci computed on a population of cells and reveal a wide range of three-dimensional chromatin structures that can be involved in various biological processes, such as DNA replication and segregation (3), gene regulation (4) or cell differentiation (5). Unusual genome organization can also be observed in disrupted biological functions such as cancer in humans (6) or infection (7).

To address the growing amount of Hi-C data produced by the community, numerous analysis tools and pipelines have been proposed covering a variety of functions, notably to process, filter and normalise (8), detect loops (9), topological domains (9), quantify compartment signals (10) or improve the resolution of poorly covered data with deep learning (8). As with other types of next-generation sequencing (NGS) and associated downstream analysis pipelines, the size of the generated reads, which is limited, for example, to the order of 50 bp, can lead to ambiguous mapping within the genome during the alignment process. Consequently, Hi-C analysis pipelines are limited to the exploitation of read pairs in which both partners of the pair align to unique genomic positions (uniquely mapping reads). Read pairs in which at least one partner is mapped at several positions (multi-mapping reads) are traditionally filtered out, leading to a significant loss of information on the spatial organization of the genome. These portions of the genome with repeated sequences remain invisible, appearing as empty bins, i.e., white lines and colons on Hi-C contact maps, representing approximately 15% of the standard Hi-C library for yeast and up to 35% of the Hi-C libraries for humans (**Supplementary** Figure 1).

However, various analyses have shown that repeated elements could play a fundamental role in metazoan genome folding (11), in the genome organization of filamentous fungi (12), in the immune escape of apicomplexan parasites such as *Plasmodium falciparum* (13), or in super-integron structure in *Vibrio cholerae* (14). One interesting recent example is the case of primate-specific endogenous retrotransposon human endogenous retrovirus subfamily H (HERV-H), which has recently been highlighted in the formation of topologically associating domains (TADs) (15). Hence, characterizing the role of repeated elements in the three-dimensional structure of genomes remains a challenge that needs to be investigated.

To our knowledge, very few pipelines have been proposed to offer a reasonable solution for processing reads with ambiguous alignments in Hi-C or MicroC experiments, and to date, there is only one pipeline addressing this issue called mHiC (16), which reallocates some of the ambiguous reads using some paired-end Hi-C read features. This pipeline allows fractional contact (i.e., multiple genomic coordinates can be assigned for one given Hi-C read pair), which can make signals difficult to interpret. This proposed answer remains partial, leaving out a significant proportion of reads that are not reallocated.

Here, we propose *Hicberg*, a computational approach that enables Hi-C data processing and maximizes the rescue of multi-mapping reads through a statistical inference based on the computation of statistical trends derived from both polymer physics and the technical features of Hi-C protocols. This approach paves the way for a deeper characterization and understanding of repeated elements in the context of the spatial organization of genomes. Although initially developed for processing data from 3C-derived capture techniques, Hicberg can also be used to process other paired-end omics, such as Mnase-seq or ChIP-seq. This allows contact signals and other genomic signals to be confronted in the same computational framework, allowing a more exhaustive exploration of signals from repeated elements.

## Results

### The algorithm’s operating principle of Hicberg

The operating principle of the computational method we propose here is to compute and use statistical trends present in the uniquely-mapping part of the data to determine the probability of the different genomic positions of possible alignments of multimapping reads (**Figure 1**).

**Figure 1.**
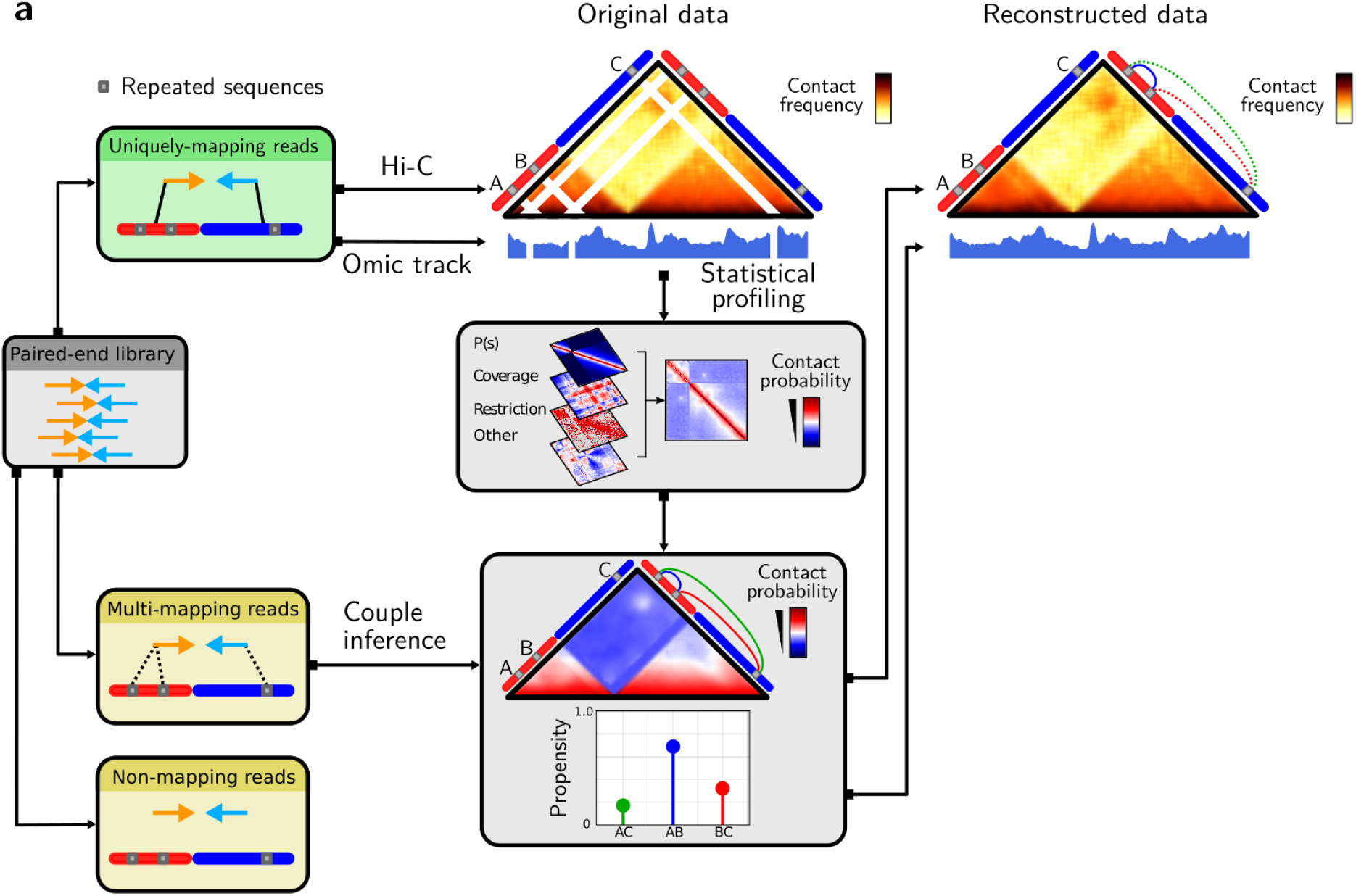
General principles of the Hicberg method for the reconstruction of contact maps and genomic tracks **a** Read pairs reconstruction involves classification into 3 groups (non-mapping, uniquely-mapping and multi-mapping reads). The uniquely-mapping fraction of reads is used to profile the statistical trends of the data. Several components can be included in the statistical model: chromatin polymeric behavior P(s), genome coverage, distances to the closest restriction site and others. The chosen user components are combined to provide the statistical model, which computes the propensities for each potential contact to occur, and the propensity distribution is built. For each ambiguous pair, a single possibility is then chosen according to the propensity distribution.

Hicberg used paired-end next-generation sequencing (NGS) data such as the Hi-C library and a reference genome as inputs. Each of the 2 mates is independently aligned to the reference genome via mapper software such as Bowtie2 (17). The alignment procedure is performed so that all possible alignment positions for an ambiguous read are given. In the case of complex genomes rich in repeated elements, a partial report of the different alignment possibilities for an ambiguous read (e.g., 100 possible alignments) can be used to limit the use of computational resources.

Hicberg then classified pairs of reads into 3 different groups (**Figure 1, Supplementary Figure S1b**): non-mapping pairs, uniquely-mapping pairs, and multi-mapping pairs. Non-mapping pairs have at least one read that does not align to the reference genome (which could be due to contamination, errors from sequencing, the presence of adapters, chimeric reads, etc.) and are excluded. Uniquely-mapping pairs, where both reads uniquely align on the reference genome. These pairs are retained and used to build the statistical model. Multi-mapping pairs have at least one mate with more than one possible alignment, i.e., the alignment quality is below a user-defined quality threshold (by default mapq > 35). The pairs of reads belonging to this last group are those that are reassigned via the statistical model.

Statistical trends are computed via the fraction of uniquely-mapping read pairs. The statistical model can rely on several tendencies present in the data. First, chromosomes can be characterized by their polymer behavior (18). Constrained by thermodynamics, in the case of intrachromosomal contacts, two loci are more likely to interact at short distances than at long distances. This observation is known as the distance law or P(s), which represents the probability (or normalized frequency) that two loci separated by distance s are in contact (19). The distance law can be split into 3 different sublaws, taking into account the directionalities of the aligned reads and reflecting the different possible read configurations associated with the Hi-C protocol (20) **(Methods, Supplementary Figure S1c)**. The signal distributions of these different configurations are different and are thus exploited within the statistical model. This point represents a significant improvement in comparison with earlier attempts to reconstruct contact signals (16). Indeed, Hicberg’s aim is to reconstruct all the signals present in a Hi-C library, whether they are informative events or those representing byproducts of the protocol. This strategy enables a more faithful reconstruction of the entire dataset.

Second, genome coverage (number of reads at each bin along the genome) is not necessarily uniform **(Supplementary Figure S1a)**. Therefore, the greater the coverage of a genomic region is, the greater the probability that a read will come from that region. This statistical trend is particularly discriminating when it presents a particular profile, such as for bacteria undergoing rapid growth, where the region close to the origin of replication is naturally 2--4 times more covered than the region close to the replication terminus **(Supplementary Figure S1a)**. Such variation in genomic coverage can thus be used as a selective component for the relocation of multi-mapping pairs.

Finally, Hi-C reads are enriched in the vicinity of restriction sites used in the Hi-C protocol (21). Thus, the closer a Hi-C read is to a specific restriction site, the more likely it is to belong to the corresponding locus. We derived a third component to our statistical model on the basis of the distribution of reads according to their distance to the closest restriction site, which is the sum of the distances of each mate **(Supplementary Figure S1a).** This parameter is noted D_1_D_2_ distance and is particularly useful for Hi-C libraries in which a single restriction enzyme is used during the digestion step.

As a multicomponent statistical model, each level of information, P(s), coverage and D_1_D_2,_ is considered independent. The model can be customized by the user to select one, several or all the components to be used for genomic coordinate reassignment of multi-mapping reads. Once the statistical model has been established, all possible pairs resulting from reads corresponding to repeated sequences are evaluated. Each of the descriptors outlined above are evaluated and combined. In this way, each plausible pair of reads is assigned a *propensity*, forming a discrete probability distribution for all possible pairs. One pair of reads is then chosen by a random draw according to this probability distribution. Ambiguous read pairs whose genomic coordinates have been inferred are then aggregated with uniquely mapping pairs, enabling the establishment of complete genomic datasets.

## Benchmark of Hicberg

We first assessed the quality of the reconstructions proposed by Hicberg via data-driven simulations based on in silico modification of Hi-C read alignment output files (“Methods” and **Figure 2a**). The benchmark strategy is based on a controlled injection of ambiguity into the alignment procedure: for certain regions, possible alignments are given as they are coming from repeated elements to generate an alternative alignment output file on which Hicberg is applied. The genomic data are reconstructed and compared with the ground truth via Pearson correlation. We also compare reconstructions from Hicberg and mHiC, the only similar tool available for Hi-C contact map reconstruction.

**Figure 2.**
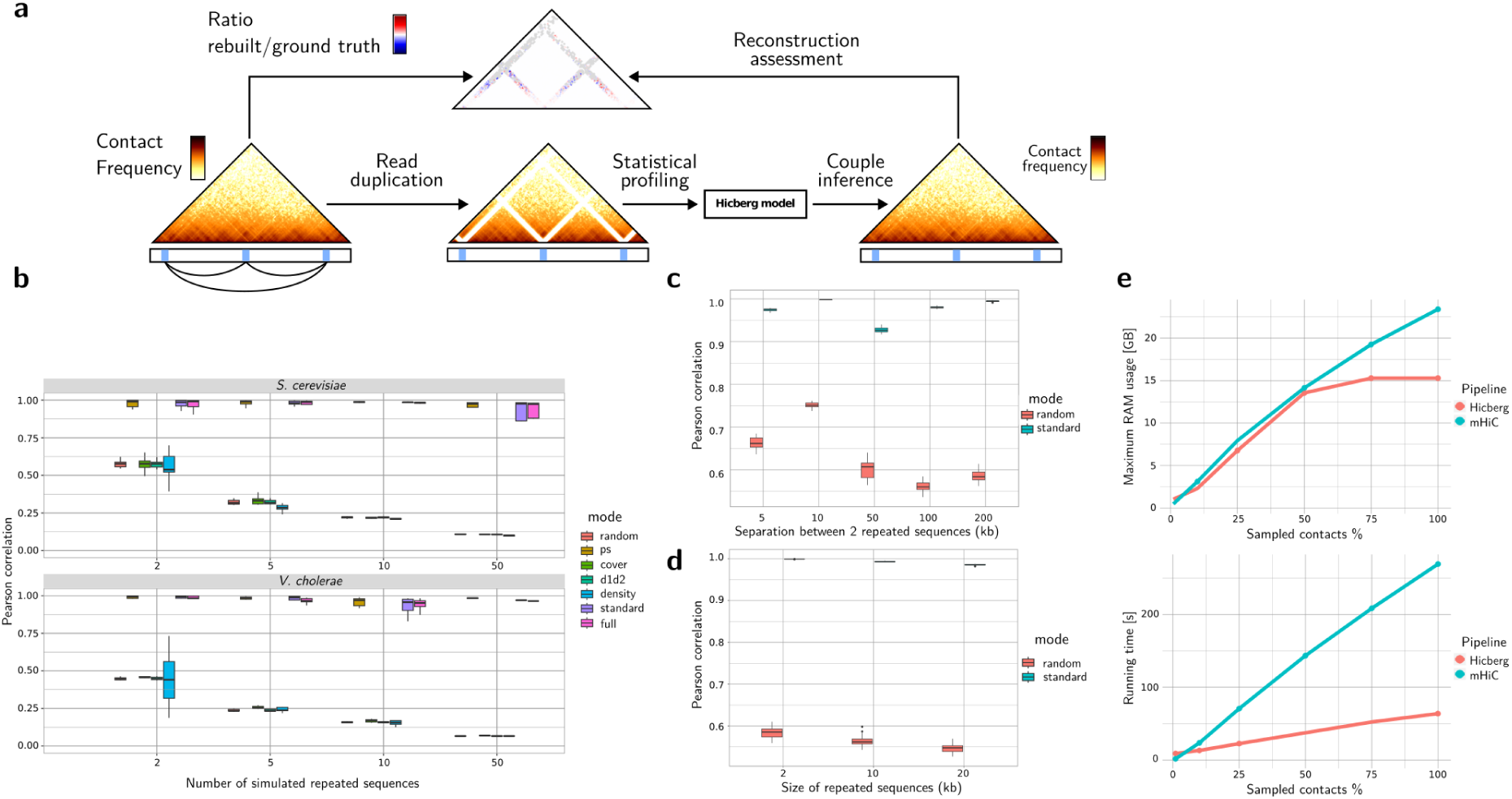
In silico bench testing strategy for assessing the quality of Hicberg reconstructions **a** Controlled ambiguity is injected by duplicating selected reads in the alignment files at alternative positions, mimicking the alignments obtained during alignment on repeated sequences. This ambiguity is then confronted with Hicberg reconstruction after profiling and statistical inference. The ground truth and the data reconstructed through Hicberg are compared to assess the quality of the proposed reconstruction. **b** Evaluation of the Hicberg’s reconstruction quality (Pearson correlation) as a function of the number of repeated sequences. For each number of repeated sequences, all statistical modes of Hicberg were evaluated on the same dataset. The top figure shows the evaluation performed on an asynchronous *S. cerevisiae* Hi-C library (47 million reads). The bottom panel presents the results of the evaluation performed on a Hi-C library of *V. cholerae* in the exponential phase (114 million reads) from (56). **c** Evaluation of the Hicberg’s reconstruction quality (Pearson correlation) as a function of the genomic distance separation between 2 repeated sequences. **d** Evaluation of the Hicberg’s reconstruction quality (Pearson correlation) as a function of the size of 2 repeated sequences separated by 30 kb. For each assay, Hicberg reconstructions were assessed in both random and standard modes (P(s) and coverage components). Assessments were performed on an asynchronous *S. cerevisiae* Hi-C library. **e,** Memory usage of both Hicberg and mHiC according to the subsampling of Hi-C data of *S. cerevisiae* from 5% to 100% of the 47 million pairs of reads of the data. Runtime of both Hicberg and mHiC according to the subsampling of Hi-C data of *S. cerevisiae* from 5% to 100% of the 47 million pairs of reads in the data.

Given the exhaustive nature of the scenarios that could be tested, we have chosen to limit ourselves to evaluations close to biological scenarios. The first parameter that can be tested concerns the number of repeated sequences within a genome. Hicberg shows strong stability to the number of repeated sequences, with a reconstruction score above 0.9 for a number of 50 when used in the standard mode (which takes into account P(s) and coverage) or full mode in *Saccharomyces cerevisiae*. The correlation coefficient decreases sharply when a random model is used or when P(s) is not considered, reaching approximately 0.12 under these conditions. The same behavior can be observed with *Vibrio cholerae* Hi-C data, with a reconstruction score higher than 0.9 when standard or P(s)-adjusted models are used (**Figure 2b**).

We also measured the quality of the reconstructions proposed by Hicberg as a function of the distance separating 2 repeated elements in *Saccharomyces cerevisiae*. We observed relative stability, with reconstruction quality scores above 0.9 for separations between repeated sequences increasing from 5 to 200 kb in standard mode, whereas scores between 0.55 and 0.75 on average were obtained for the same separations in random mode. The dispersion of the reconstruction quality scores is also greater when a random model is used.

Finally, we evaluated the quality of the reconstructions according to the size of the repeated sequences. Hicberg shows robustness with respect to this parameter, with reconstruction qualities over 0.97 up to 20 kb in *Saccharomyces cerevisiae* in standard mode, compared with scores of less than 0.6 with a random model (**Figure 2c**).

For comparison with computer resources with the only tool already available, we ran Hicberg and mHiC on standard Hi-C data from an asynchronous *S. cerevisiae* yeast population (Methods) with a sequencing depth of 47 million reads. Hicberg outperforms mHiC in terms of both RAM usage and computing time. The Hicberg RAM consumption level remains well below mHiC regardless of the level of data subsampling (**Figure 2d**). In terms of computation time, when 16 CPUs are used, contact map reconstruction with 2 kb resolution according to standard parameters is achieved in 63 minutes with Hicberg versus 270 minutes for mHiC from read alignment to contact map construction without library subsampling (**Figure 2d**).

A major difference between Hicberg and mHiC is the tolerance of contact splitting for a pair of reads to be reassigned (i.e., the possibility of assigning several genomic coordinates to a given pair of reads). Our choice with Hicberg not to tolerate this splitting makes absolute comparison of genomic track reconstructions via our strategy complex. Indeed, using the standard mHiC parameters, we obtain matrices with a greater overall number of contacts than the number of initial reads in the library under study. Although not quantitatively evaluable according to the modality set for the Hicberg benchmark, the quality of the reconstructions proposed by our method and mHiC can be visually appreciated. Indeed, the reconstructions proposed by mHiC of the subtelomeric regions of chromosomes IX and X in *Saccharomyces cerevisiae* (**Supplementary Figure S3 a and b**) show an absence of contact reattribution and therefore an absence of correctly defined structuring of these regions, in contrast to the structuring proposed by Hicberg. A similar observation can be made for reconstructions of the peripheral region of rDNA located on chromosome XII in *Saccharomyces cerevisiae*. This region, previously absent from contact maps, is very poorly reconstructed via mHiC, whereas a clearly defined structure, showing local chromatin compaction, was revealed by Hicberg (**Supplementary Figure S3 c**). Finally, some specific regions, such as those located on chromosomes VII and XIV, presented medium-range artifactual patterns after mHiC reconstruction, even though these patterns were absent from Hicberg reconstructions (**Supplementary Figure S3 d and e)**.

### Reconstruction of genomic data in *Saccharomyces cerevisiae*

To illustrate examples of new biological analyses that can be performed via Hicberg reconstructions, we first focus on reconstructions of chromosome contact maps in the model organism *Saccharomyces cerevisiae*. This yeast has a medium-sized genome of 12.5 Mb, with diverse sources of repeated sequences: it has five LTR-retrotransposon families called Ty1, Ty2, Ty3, Ty4 and Ty5 (22), with approximately 50 elements distributed across all chromosomes. It contains the rDNA (ribosomal DNA) region, which consists of a 100 to 150--copy locus repeat. It also contains repeated sequences in telomeric regions, which mainly contain two types of repetitive elements, called “X” and “Y’”. Finally, it contains several duplicated genes (e.g., *CUP1* on chromosome VIII). For alignment, we used the genome sequences recently obtained for telomere-to-telomere (T2T) quality (23,24) associated with the online database ScRAPdb (see “Methods”).

We started performing Hicberg reconstructions on an example of standard Hi-C libraries (using a commercial kit, Methods) for the yeast *S. cerevisiae* under normal growth conditions and for an asynchronous population. Without Hicberg reconstruction and with standard filtering of ambiguous reads, 35.3 million exploitable reads remained over the initial 47 million. With Hicberg reconstruction, there are 42.3 million exploitable reads, representing 7 million additional pairs of reads, i.e., an information gain of 20%. The remaining 5 million read pairs where at least one of the 2 reads is chimeric or unmappable owing to sequencing errors or mutations, sequencer noise, etc.

We present an example of chromosome VII reconstruction (**Figure 3**) as well as for all chromosomes (**Supplementary Figures S4 to S21**). The coverage of the reconstructed parts reaches a normal level for the whole chromosome, and the contact map shows a homogeneous signal on all the different chromosomes. With respect to reconstruction, some regions still have very low coverage. This is the case for chromosome I, for example, where several bins show low Hi-C coverage with reconstruction (**Supplementary Figure S4**). Gene annotation revealed that these genes correspond in particular to those of the FLO family (e.g., *FLO9* and *FLO1* for the chromosome I). Computation of the number of restriction sites used in the Hi-C protocol (DpnII, HinfI) revealed that all FLO genes were deficient in these restriction sites (with the exception of the *FLO8* gene on chromosome V) (**Supplementary Figure S21**). On the basis of Hi-C data from a similar type of protocol, it was proposed (25,26) that these genes present contact enrichment. The fact that these genes are restriction site deficient has a significant impact on their detectability, and different protocols are needed to study contacts between FLO genes. This highlights the fact that Hicberg can uncover regions that are difficult to detect apart from mapping problems.

**Figure 3.**
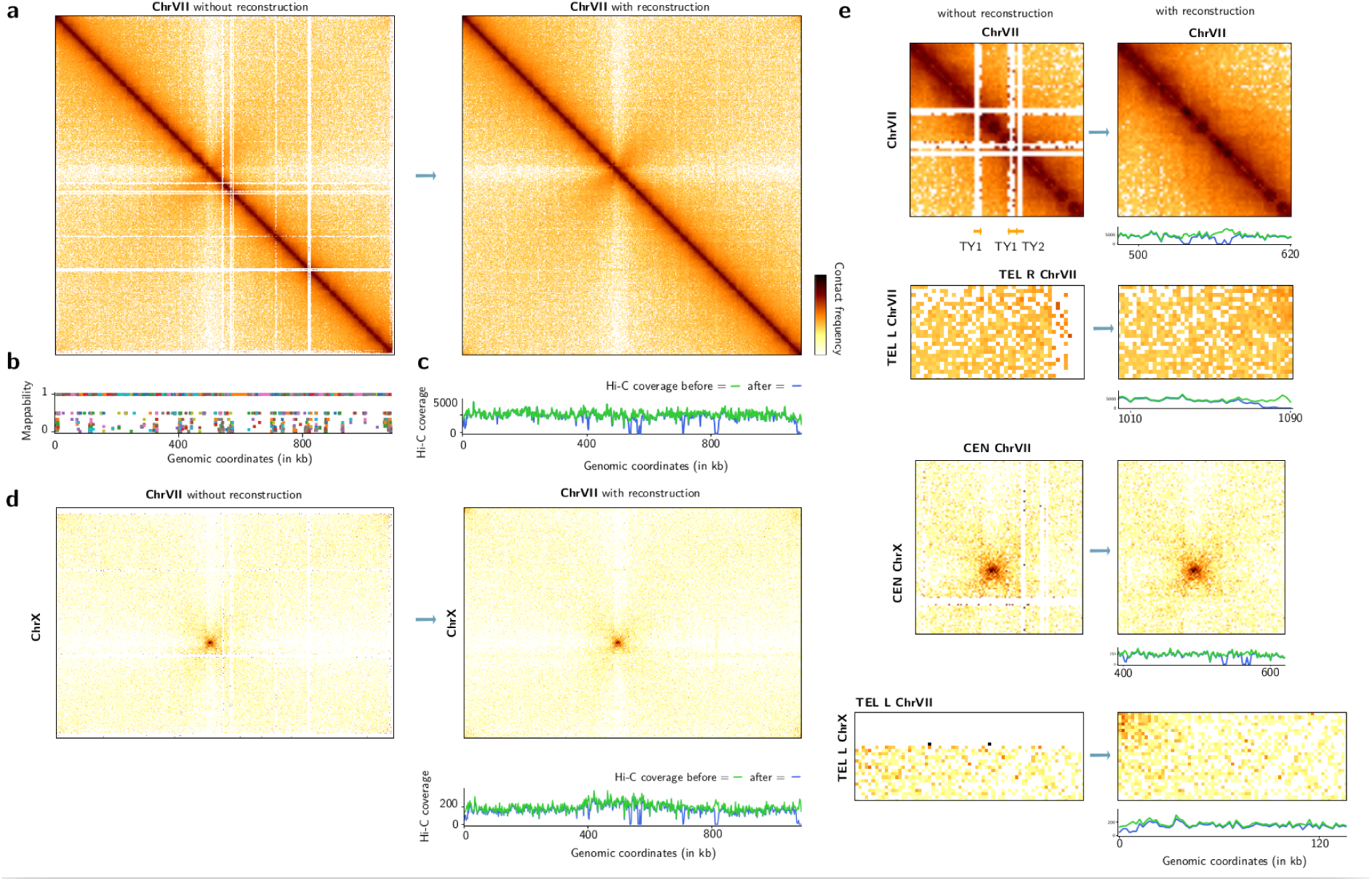
Examples of Hicberg reconstructions in the genome of the model organism *Saccharomyces cerevisiae*. **a** Chromosome contact map without and with Hicberg reconstruction for chromosome VII (bin = 2 kb, with normalisation) **b** Mappability signal along the chromosome VII. **c** Hi-C coverage before (green) and after (blue) Hicberg reconstruction. **d** Interchromosomal contact map between chrVII and chrX without and with Hicberg reconstruction (bin = 2 kb, with normalisation) with Hi-C coverage without and with Hicberg reconstruction. **e** Examples of reconstructions at representative regions (zooms of a and d): on several yeast transposons (Ty) at chrVII, between both telomeres TEL-L and TEL-R of chrVII, between centromeres of chrVII and chrX, and between telomeres TEL-L of chrVII and TEL-L of chrX with Hi-C coverages without and with reconstruction.

Hicberg reconstructs specific contact signals between telomeres on the same chromosome or between different chromosomes (**Figure 3 e**). These signals are important biologically, as they depend on the cell state and metabolism. For example, in quiescent cells, the grouping of telomeres into a unique focus or hypercluster localized in the center of the nucleus has been shown (27,26).

Chromosome reconstructions revealed interesting contact patterns, such as the presence of chromosomal domains at subtelomeric regions such as chromosome IX (**Supplementary Figure S11 b)** or chromosome XIII (**Supplementary Figure S15 b).** We also observed a border pattern at the HML locus (Mating type cassette - left, the mating type cassette - left) on chromosome III (**Supplementary Figures S6 b),** which recalls the border pattern at the HMR locus visible in the data without reconstruction (**Supplementary Figures S6 b).** Interestingly, these loci have been recently proposed to be associated with loop extrusion processes with condensin complexes (28).

We also present 2 examples of contact maps between 2 different chromosomes (**Supplementary Figure S20**). Reconstructions can show lines in depleted areas of contact (notably at centromeres). However, these reads represent a very small proportion of the signal.

Finally, we generated a mitochondrial contact map (**Supplementary Figure S19**). Although reconstruction with Hicberg only slightly increases map coverage, it is possible to see an interaction domain at the level of the large *COX1* gene (size 12.8 kb), reminiscent of domain structures observed in bacterial genomes (29).

## Hicberg can detect changes in repeated sequence positions in a reference genome

Aberrant contact patterns can appear in several contact maps, for example, at chromosome III and chromosome XII, with reconstructions using the current reference genome of the *Saccharomyces* Genome Database (SGD) (Genome assembly R64, GCF_000146045.2, sacCer3) instead of the T2T assembly (**Supplementary Figure S22 a)**. By plotting the 4C-like contact signal as, for example, from a family of Ty1 retrotransposons (Methods), it is possible to detect peaks at regions that are not annotated as carrying a Ty (**Supplementary Figure S22 b,c)**. Indeed, this type of approach allowed the identification of the presence of a Ty1 on chromosome XII not annotated in the Saccer3 version of the *S. cerevisiae* S288C genome. A PCR experiment followed by Sanger sequencing confirmed the presence of Ty1 at this genomic position (**Supplementary Figure S24 d, Methods)**. This finding shows that the Hicberg approach can be an alternative method to long-read sequencing for the discovery, annotation or confirmation of transposons in diverse genomes.

## Reconstruction of genomic data in bacteria

Finally, we present 2 examples from the bacterial world: the first example corresponds to the reconstruction of a contact map of the genome of the bacterium *Escherichia coli*, which contains a duplicated region of several kb (**Supplementary Figure S23)**. The reconstruction shows a map consistent with the layout of the duplication, enabling us to check the geometry and correct positioning of genetic constructs within the genome involving repetitions. We also present an example of a reconstruction for the *Vibrio cholerae* bacterium, which contains 2 chromosomes (**Supplementary Figure S24)**. The reconstruction is able to produce a contact pattern between the 2 chromosomes, notably at the level of the superintegron on chromosome 2, which harbors hundreds of gene cassettes (30).

## Hicberg is able to reconstruct signals linked to compartmental organisation

To explore the potential of Hicberg reconstructions with complex contact signals, we reconstructed several Hi-C datasets of organisms that have compartment signals. This characteristic of the spatial organisation of genomes, particularly in eukaryotes, notably in humans (2), corresponds to a bipartitioning of the genome into 2 compartments: one, known as compartment A, which is rather active at the transcriptional level and contains genes, and another, known as compartment B, which is rather silent and contains fewer genes. This signal is particularly visible on the correlation matrix in the form of a chequered pattern (31). Our laboratory has recently artificially created a compartment signal in chimeric *S. cerevisiae* strains that contain pieces of the *Mycoplasma mycoides* bacterial chromosome on chromosome XVI (32). This chromosome alternates between a yeast chromosome with a GC content of approximately 38% and an *M. mycoides* chromosome with a GC content of approximately 20% (**Figure 4a**).

**Figure 4.**
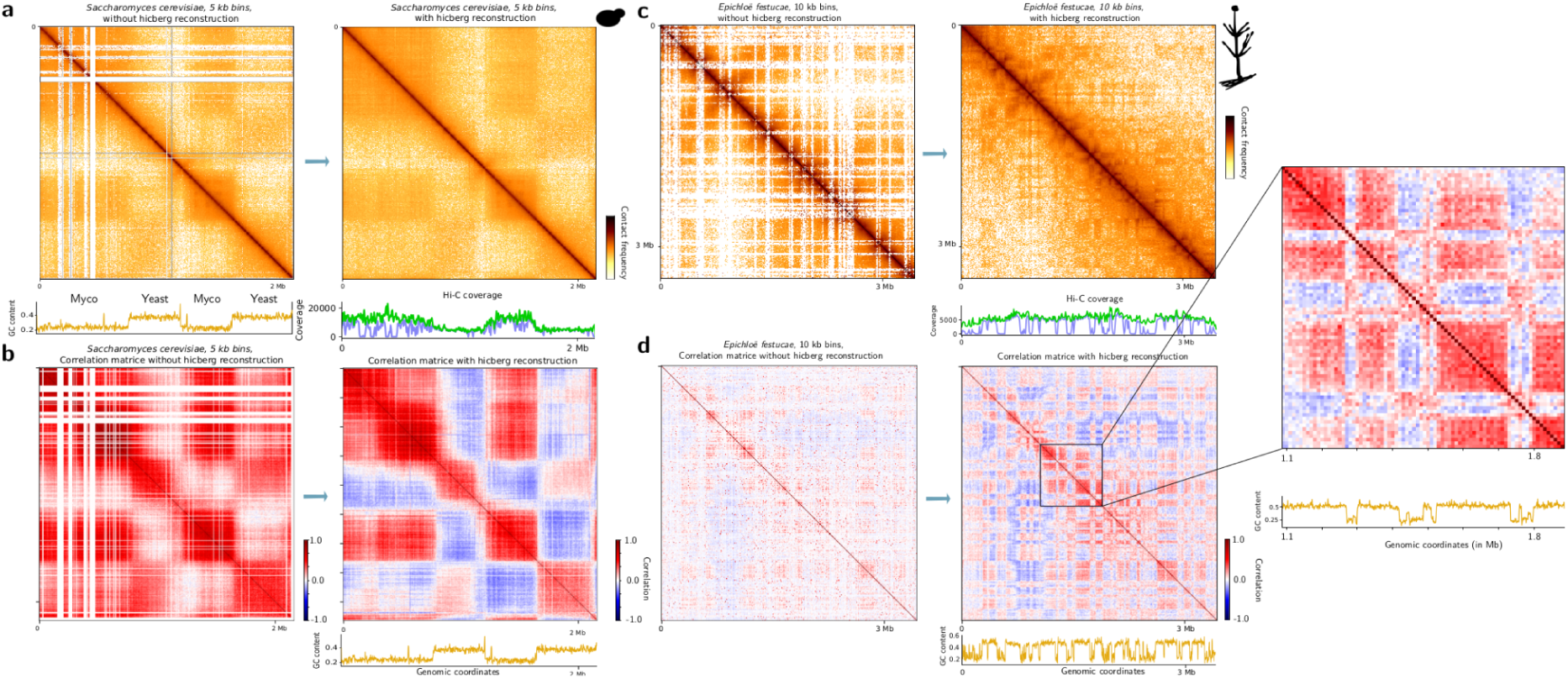
Compartment reconstruction signals by Hicberg **a** Normalised contact map of the chimeric chromosome XVI chromosome of *S. cerevisiae* and *M. mycoides* binned at 5 kb resolution without and with Hicberg reconstruction. The GC content of the sequences and annotations for *S. cerevisiae* and *M. mycoides* are shown below the left, and the Hi-C coverages before (blue) and after (green) the Hicberg reconstructions are shown below the right. **b** Correlation matrices of the chimeric chromosome XVI of *S. cerevisiae* and *M. mycoides* binned at 5 kb resolution without and with reconstruction. The GC content is shown below the map on the right. **c** Normalised contact map of chromosome 5 of *Epichloë festucae* binned at 10 kb resolution without and with reconstruction. Hi-C coverages before (blue) and after (green) Hicberg reconstruction are shown below right. **d** Correlation matrices of chromosome 5 of *Epichloë festucae* binned at 10 kb resolution without and with reconstruction. The GC content (orange) is shown below the map on the right. A zoom of the contact map is shown in relation to the GC content signal below.

*M. mycoides* bacteria contain several repeated sequences, particularly insertion sequences (ISs), which are bacterial transposons that can affect genome organisation and plasticity (33). These regions generate white lines at the IS regions in the contact maps processed with standard pipelines. With the Hicberg reconstructions, the normalised contact map shows a homogeneous signal over the whole chromosome, including the repeated sequences present on the *M. mycoides* segments (**Figure 4a)**. Interestingly, if we plot the 4C signal, such as that of the telomeric region present only in the reconstructed data, we can observe that the signal shows nonmonotonic behavior with increases in intermediary genomic distances in the contact signal following the organisation of the compartments (**Supplementary Figure S25)**.

The computation of the correlation matrix used to highlight the compartment signals is much clearer, with a stronger correlation signal across the whole matrix (**Figure 4b)**. While the Hicberg reconstructions on the normalised matrices mainly modify the lines with poor coverage, the effect of the Hicberg reconstructions on the correlation matrices, which calculate for each pixel the correlation coefficient between the whole row and the whole column, has an effect on the whole correlation matrix.

We then tested it on other microorganisms naturally containing a high proportion of repeated sequences. For example, we tested it on the organism *Epichloë festucae*, which is a filamentous fungus known to have a high proportion of repeated sequences in its genome, notably LTR retrotransposons, Miniature Inverted Repeat Transposable Elements (MITEs) (34) and DNA transposons. It was shown that repetitive elements play a key role in the nuclear organisation of *E. festucae*. These repetitive elements help to divide the genome into distinct regions with similar gene expression profiles (12). Without reconstruction, a large proportion of reads are below the mapping threshold of standard pipelines, generating many empty lines or presenting an unusable weak contact signal (**Figure 4c**). The Hi-C coverage signal shows a very inhomogeneous profile with many undetectable regions. The correlation matrix used to highlight the compartment signals is also poor in exploitable signals, and only a restricted part of the genome allows us to observe a checkerboard signal typical of the behavior of the signal in the compartments (**Figure 4d**). With Hicberg reconstruction, the contact map is filled with signals, the Hi-C coverage becomes homogeneous, and the compartment signal can now be observed at the scale of the chromosome and the whole genome. Interestingly, this confirms the link with GC content: genomic regions with similar GC contents are present in the same compartment (**Figure 4d**, zoom).

Both of these examples of reconstructions display Hicberg’s ability to reconstruct signals that deviate from the average behavior mediated by contact frequency as a function of genomic distance (P(s)). Indeed, the reads present in the dataset provide sufficient constraints for signal prediction and allow the reconstruction of more complex signals for compartment signals.

## Parasitic 2µ plasmid present enrichment of contact with Ty transposons

The 2µ plasmid is a natural plasmid present in the nucleus of laboratory or natural strains of *S. cerevisiae* (35). It is 6.4 kb in size and does not appear to confer any advantage on its host (36). It has very high stability, close to that of its host chromosomes, potentially due to a hitchhiking mechanism (37). Recently, the 2µ plasmid was shown to preferentially contact long regions with low transcriptional activity (38). Compared with the average size of genes (∼1 kb) of the host organism, Ty retrotransposons in the yeast *S. cerevisiae* are also rather large (39), with the size of approximately 5.5 kb. The transcription of these genes is generally low compared with that of the rest of the genome (40). They meet the 2 criteria that make them potential hot spots of contact for the 2µ plasmid. To test this hypothesis, we reconstructed the contact signals of the 2µ plasmid with the different elements making up the Ty1, Ty2, and Ty4 families. The contact signal of the 2µ plasmid is enriched with various Ty elements (**Figure 5 a–d**).

**Figure 5.**
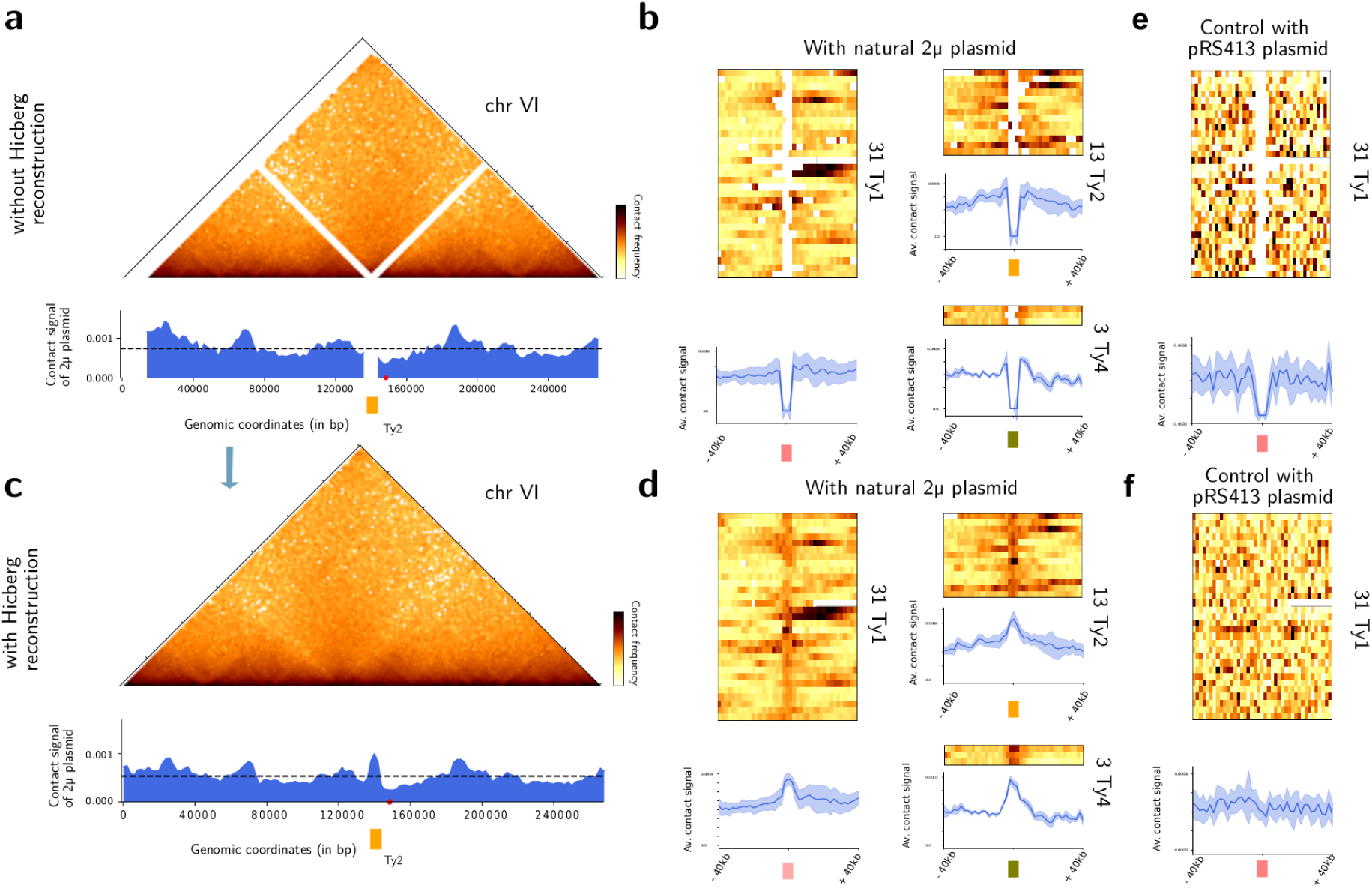
Parasitic 2µ plasmid present enrichment of contact with yeast transposons **a** Contact map of chromosome VI of *S. cerevisiae* without reconstruction; bin = 2 kb in the asynchronous population (BY4741 strain). Below the contact signal of the 2µ plasmid along chromosome VI without reconstruction. **b** Heatmap representing the contact signals of the 2µ plasmid with 3 different families of yeast transposons: Ty1, Ty2 and Ty4 without reconstruction. Below the averaged signal (in blue) over the different elements of each family. **c** Contact map of chromosome VI of *S. cerevisiae* with reconstruction, bin = 2 kb. Below the contact signal of the 2µ plasmid along chromosome VI with reconstruction. **d** Heatmap representing the contact signals of the 2µ plasmid with 3 different families of yeast transposons: Ty1, Ty2 and Ty4 with reconstruction. Below, the averaged signal summed over the different elements of each family. **e** Heatmap representing the contact signals of the pRS413 plasmid sequence with Ty1 elements without reconstruction. Below, the averaged signal over the different elements for the Ty1 family. **f** Heatmap representing the contact signals of the pRS413 plasmid sequence with Ty1 elements with reconstruction. Below the averaged signal summed over the different elements of the Ty1 family.

The average contact profile for each of the families shows contact enrichment. For the negative control, the same analysis was performed with the pRS413 plasmid, which does not have the 2µ gluing system but relies on a centromeric system for its partitioning mechanism, and the contact enrichment is not present (**Figure 5 e**). This is also the case for other sequences used as negative controls: the mitochondria or other chromosomes of the host do not show enrichment with these same families of retrotransposons (data not shown). Interestingly, these reconstructions show that these two biological objects, which are in a sense two colonizers of the *S. cerevisiae* genome, colocate inside the nucleus.

## Ty transposons can participate in the positioning of cohesins

Chromosomal loops have recently been shown to form at certain stages of the cell cycle in mammals (41) as well as in the model organism *S. cerevisiae* (42). In the yeast *S. cerevisiae,* loops of 10–15 kb are formed by cohesin complexes in the G2/M phase (43,44), and their size can be finely regulated by a set of dedicated proteins, such as Pds5 or Wapl (45). Cohesin complexes, which are molecular motors, can condense the whole genome, possibly through loop extrusion (46), which can impact various biological processes, such as DNA repair (47). The minimum ingredients for the formation and precise positioning of these loops are still under debate. Some models, for example, have recently proposed that transcription is the source of energy that drives cohesin and engages it in chromosomal loops (48). To explore the potential impact of the presence of retroposons on cohesin loop positioning, we produced reconstructions of chromosome Hi-C contact maps under 2 different biological conditions: in the G1 phase, cohesins are not bound to chromatin, and in the G2/M phase, the cdc20 genetic mutant, which synchronizes cells in the G2/M phase, was used. The data used were extracted from (47) in the W303 genetic background with the T2T assembly genome. Interestingly, we observed that retrotransposons placed in a convergent configuration, such as two pairs on chromosome IV, are associated with the presence of chromosomal loops only in the G2/M phase (**Figure 6 a, b, c, d;** black arrows). Interestingly, the genome of strain W303 contains a third pair of retrotransposons in a convergent configuration, which also shows a loop pattern at the convergence area of Ty (**Supplementary Figure S26**).

**Figure 6.**
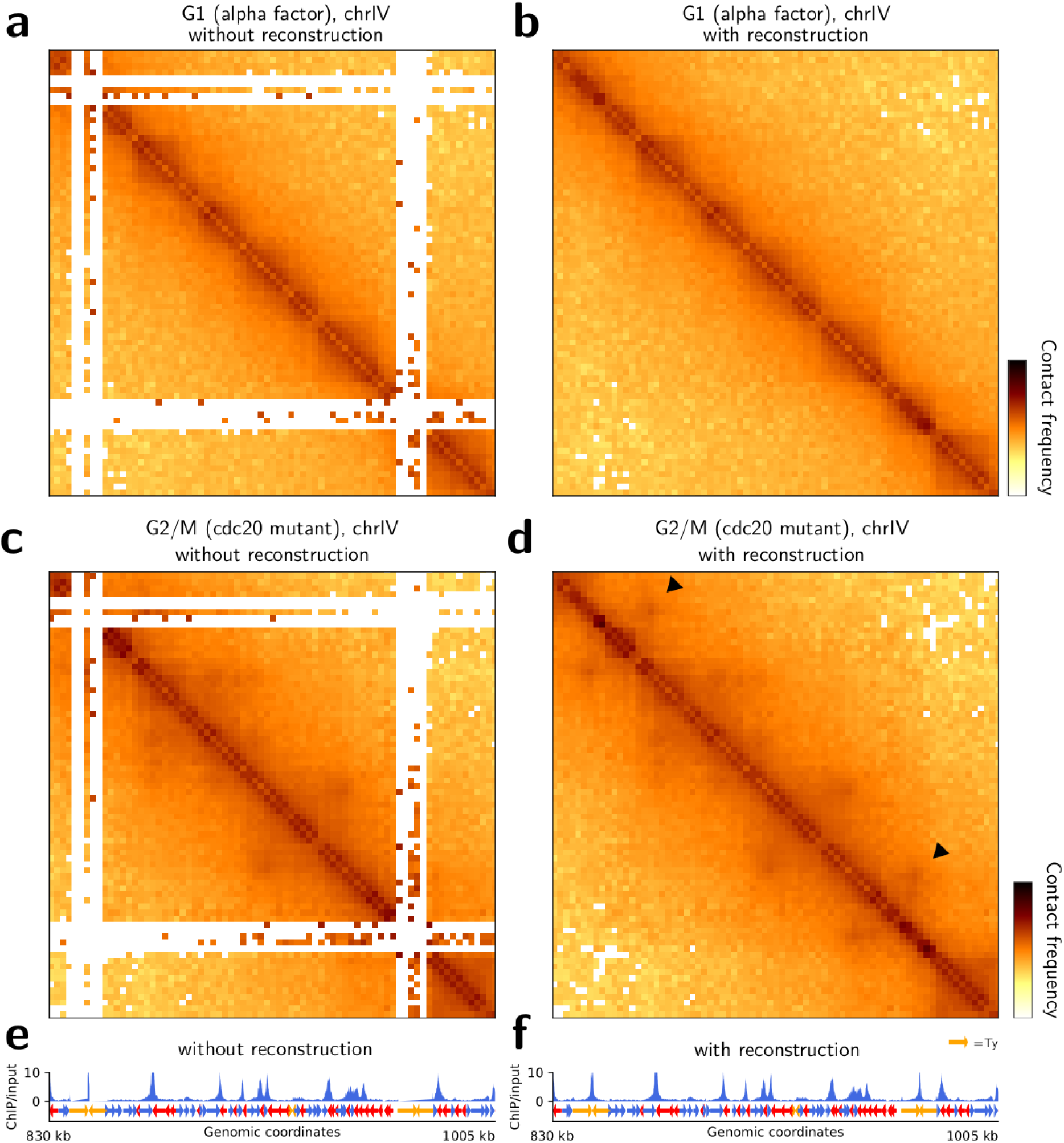
Yeast transposons can participate in the positioning of cohesin in the *S. cerevisiae* genome **a–b** Zoomed view of the contact map of chromosome IV of *S. cerevisiae* without and with Hicberg reconstruction; bin = 2 kb in a population synchronized in the G1 phase (W303 strain, data from (47)). **c–d** Zoomed contact map of chromosome IV of *S. cerevisiae* without and with Hicberg reconstruction in a population synchronized in the G2/M phase (W303 strain, cdc20 mutant; data from (47)). **e–f** Cohesin occupancy signal from ChIP-seq of Scc1 in the G2/M phase without and with Hicberg reconstruction. The forward and reverse genes are represented by blue and red arrows, respectively. Retrotransposons are represented by orange arrows. The black triangles point to chromosomal loops that are only visible with Hicberg reconstruction and are associated with cohesin occupancy peaks (Scc1) at convergent transposons.

Hicberg can also be used to reconstruct 1D tracks of various genomic signals, such as nucleosome occupancy by Mnase-seq, protein occupancy by ChIP-seq or chromatin accessibility by ATAC-seq. To confront the chromosomal loops observed at the ends of Ty elements, we reconstructed the occupancy signal for the protein Scc1, which is part of the cohesin complex, and synchronised it with contact maps (**Figure 6 e, f**). Interestingly, we detected peaks of occupancy at the ends of the open reading frames (ORFs) of the Ty that form chromosomal loops. Notably, the peaks are sometimes already partially visible in the data without reconstruction.

To generalize the analysis, we constructed an agglomerated plot of the occupancy signal for the protein Scc1 for all Ty elements and for all genes of *Saccharomyces cerevisiae* without and with Hicberg reconstruction. The Ty group is enriched at the ends of the ORF which is more pronounced (a 2-fold increase) than the average of the yeast genes (**Figure 7 a,** biological replicate in **Supplementary Figure S27 c)**. When each family of retrotransposons is calculated separately (**Supplementary Figure S27 a),** this behavior is visible for the Ty1, Ty1 and Ty4 transposon families. We replicated the analysis in the S288C genetic background, where the retrotransposons are not all at the same positions, and obtained similar results (data not shown).

**Figure 7.**
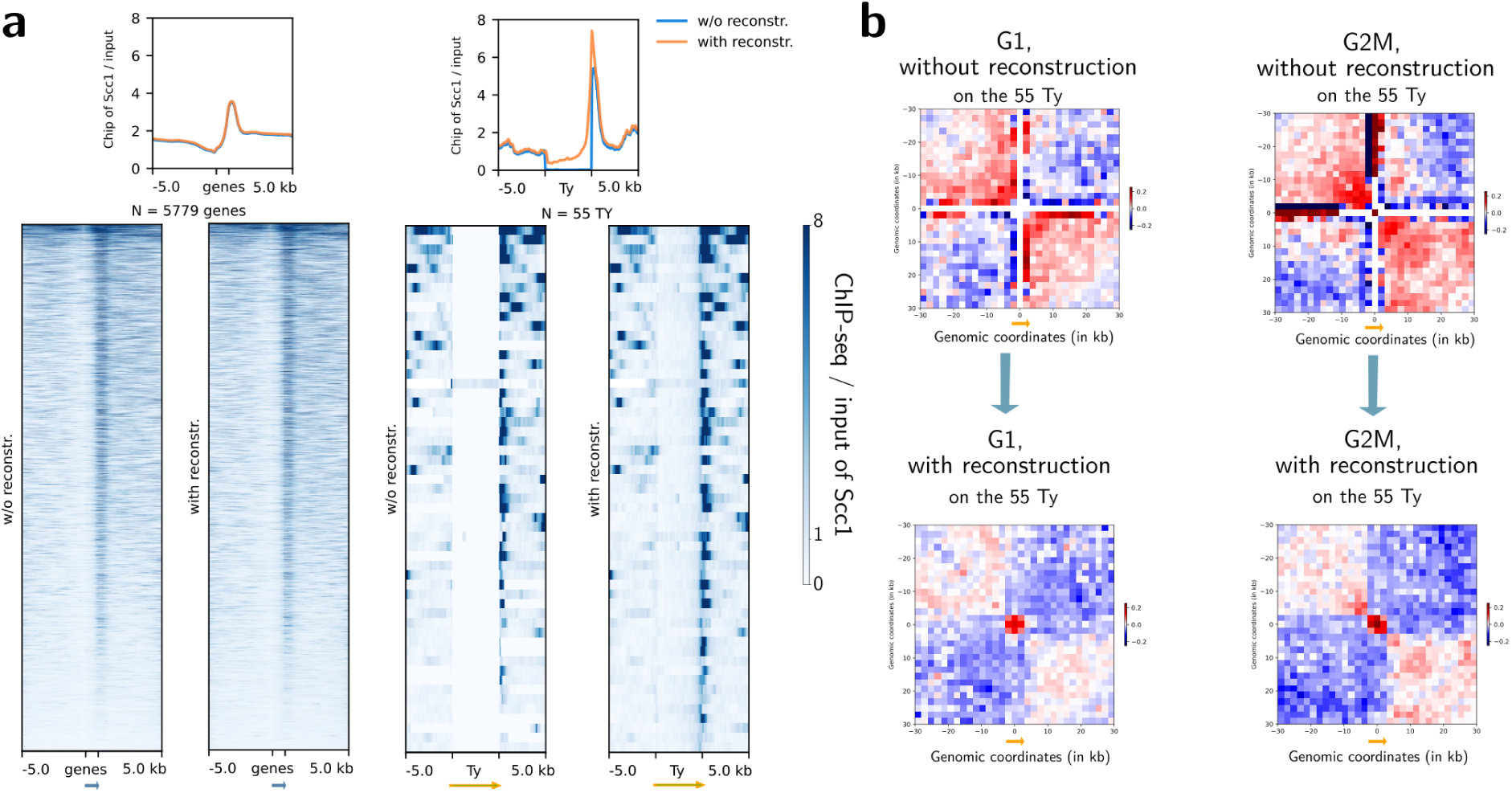
Averaged signal of cohesin occupancy and agglomerated contact plot of the transposon elements (Ty) in the yeast *S. cerevisiae* genome **a** Averaged signal of cohesin occupancy (ChIP-seq of Scc1) at genes and at Ty elements without and with Hicberg reconstruction in the yeast *S. cerevisiae* in the G2/M phases of the cell cycle. **b** Agglomerated contact plots of Ty in the G1 and G2/M phases of the cell cycle without and with Hicberg reconstruction in the yeast *S. cerevisiae*.

If we generate the average contact signal plot at the diagonal of the contact maps, we see a border pattern without and with reconstruction (**Figure 7 b)**. Without reconstruction, the average plot contains fluctuations, but the border pattern is already visible with the presence of red squares on the diagonal and blue areas around them (contact depletion). The pattern also appears to be strongest in G2/M, where cohesin is active. With only reconstruction (**Figure 7 b)**, the average plot shows a small area of the domain at the level of the Ty ORF, which has a size of approximately 5.5 kb, i.e., 3 bins, suggesting a structure linked to the Ty ORF, which is reminiscent of the gene globules observed in yeast in MicroC (49,50).

Finally, if we compute the average agglomerated signal between the different transposon elements, we do not detect any loop or enrichment pattern, indicating that the reconstructions do not show clustering of the different Ty within the nucleus (**Supplementary Figure S27 b)**.

The Ty1 retrotransposons, which have been best characterised, are not known to have high transcriptional activity (but do have high RNA stability) under normal culture conditions (39). The fact that they are associated with strong levels of cohesin occupancy at the ends of their ORF positions is an intriguing observation in the context of understanding the ingredients required for the natural cohesin positioning throughout the cell cycle. Interestingly, cohesin has recently been shown to be enriched in the 5’ region of long interspersed nuclear element-1 (LINE-1 or L1), which is a retrotransposon group that constitutes 17% of the human genome (51) and can form chromosomal loops between LINE-1 and their cognate gene promoters. All these observations suggest a particular impact of certain transposons, particularly long transposons, on the positioning of the molecular motors that organize genomes, such as cohesins.

## Discussion

In this work, we present *Hicberg*, an algorithm based on statistical inference with the computation of specific probability distributions to reassign the positions of reads from repeated sequences. To the best of our knowledge, Hicberg is the only computational method that allows the positions of multi-mapping reads to be reattributed at all spatial scales. The algorithm does not filter the data and reconstructs all the events present in the NGS library in pair-end, whether they are informative events, non-informative events or noise. This approach therefore enables the data to be reconstructed faithfully and can then be used to work on data that are more homogeneous across the whole genome. We demonstrated that our tool outperformed the only algorithm currently available for carrying out such an operation (mHiC, (16)). Hicberg paves the way for the exploration of the functional 3D organization of repeated elements. It has been designed to be versatile and usable both for all Hi-C derivative technologies and for a plethora of pair-ended omics data, as well as for a wide range of genomes and experimental conditions.

A current limitation of Hicberg reconstructions concerns regions with very low signals and high inhomogeneity, such as contact maps between different chromosomes. Our approach is sometimes unable to reconstruct the inhomogeneity of data around centromeric regions in particular. A possible extension would be to add a new level of information containing 2D signal variations, for example, via geometric extrapolations. While such a refinement is possible and is currently being explored (*density* mode in the code), it presents a significant computational overhead that does not justify its implementation in the current Hicberg package. Indeed, the fact that Hicberg is capable of reconstructing signals that deviate from the average behavior, such as chromosome loops during certain cell cycle states or compartment signals, shows that the constraints provided by the reads present in the library and the statistical inference currently implemented in Hicberg are sufficient for the reconstruction of this type of signal.

Furthermore, the use of Hicberg on contact data from strains whose genetic background could vary slightly from the reference genome resulted in the formation of unusual patterns on the contact maps, suggesting changes in the positions of repeated elements within these strains (**Supplementary Figure S22**). Since Hicberg is sensitive to variations in genetic background, e.g., positions of transposons, it can be used as an alternative tool for detecting genomic variations.

With the growing appeal of chromosome contact techniques, driven by lower sequencing costs and optimized capture methods, the Hicberg algorithm provides a bioinformatics framework for the exploration of new hypotheses concerning genome architecture and the role of repeated elements.

## Materials and methods

### Read alignment and classification

The alignment of reads to a reference genome was performed with Bowtie2 (17) in the "very-sensitive" mode. For genomes containing repeated sequences with a reasonable number of occurrences (for example, < 4,000), the *-a* option in Bowtie2 can be used, which will give all possible positions for an ambiguous alignment, which is the case for the model organisms *E. coli* and *S. cerevisiae*. For genomes with many repeated sequences, the *-k 100* option of Bowtie2 could be used, resulting in 100 possible genomic positions for a read. The SAM files are then converted to bam files via the software SAMtools.

## Computation of statistical trends

Different statistical trends are computed via uniquely-mapping pairs to create a statistical model that is used to assign ambiguous reads. Here, we briefly describe the computations of 3 of them, but others can be generated and added to the model. We assume that each of the statistical trends is independent of the others.

- The statistical law known as *P(s)* represents the contact frequency as a function of the genomic distance *s* (in bp). This metric is regularly computed in Hi-C analyses. It contains a thermodynamic component: it derives from the polymer behavior of the chromosome, and the frequency of contact decreases as the genomic distance increases. It may also contain trends due to biological chromatin condensation (e.g., during the G2/M phase of the cell cycle of the yeast *S. cerevisiae*, the appearance of regular loops every 15 kb has an impact on the behavior of this law (43)).
Finally, this law also depends on the presence of non-informative events commonly present in the Hi-C libraries, which can be classified into 3 different categories and depend on the directionality of the reads that make up the pair (20).

- “Dangling-ends” events correspond to non-digested collinear fragments that can represent a significant proportion of sequenced events in Hi-C libraries (with (+,-) orientations).
- “Self-circularized” events correspond to collinear fragments that have circularized on themselves (with (-,+) orientations).
- “Mirrors” events correspond to pairs of reads having the same orientation and aligned on the same restriction fragment (with (+,+) or (-,-) orientations).

We have therefore chosen to compute the *P(s)* law for each of these 3 possible configurations, i.e., taking into account the directionality of the 2 reads making up the pair. This allows us to take advantage of the variability in the frequencies of appearance of these different types of events, presenting a discriminating property in the attribution of genomic coordinates for the multi-mapping read pairs (**Supplementary** Figure 1a).

For contacts between different chromosomes, the average contact frequency for each pair of chromosomes is computed while accounting for the size of each of the 2 chromosomes.

- The second statistical trend we calculate is the *coverage* along the chromosomes. Each chromosome is binned at a given size (for example, 2 kb), and the number of reads is counted for each bin. A smoothing procedure is applied via a sliding window. The window size is computed on the data by taking the maximum number of consecutive bins that have significantly low values.
- The 3rd statistical trend that can be extracted from the data is the sum of the distances between the start of the reads and the next restriction site. This value corresponds roughly to the sum of the distances between the read start site and the closest restriction site and follows a specific distribution (**Supplementary** Figure 1a). The size D_1_D_2_ is computed via the restriction map of the enzymes used in the Hi-C protocol and the directionality of the reads that form the pair. The set of sizes thus detected is used to generate a characteristic distribution that constitutes a statistical trend that can be included in the model.

## Assignment of the contacts from the different possible pairs

Once these different trends have been computed, the propensity of each possible pair of reads *(i,j)* for the ambiguous alignments is computed as the product of the 3 previous statistical trends:

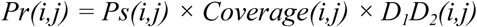

where *(i,j)* represents the genomic coordinates of a given ambiguous read pair.

The propensities of possible contacts resulting from all *(n,m)* pair combinations are computed and form a probability law:

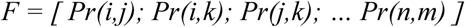

where each *Pr(i,j)* represents the weight of the probability distribution of the different possible pairs *(i,j); (i,k); (j,k) … (n,m)*. The sum of these values is normalized to 1.

The function *choice* from the *numpy.random* package that returns nonrepeated and nonuniform random samples of the given array *F.* Notably, the chosen coordinates are not necessarily those with the highest weight, which makes it possible to be more realistic about the statistical features of the data and allow for some fluctuations.

## Generation of matrices and normalisation

Pairs of contacts are then binned (by default, 2 kb bins) in contact matrices, which are normalised via the ICE method (52) and saved in the cool format (53).

## Hicberg benchmarking

To quantify the quality of the reconstructions generated by Hicberg, we adopted a simulation strategy in which we controllably inject several possible alignments for a given genomic region. The first step is to define a set of genomic intervals whose number, genomic distance or size can vary. Second, the set of (initially unambiguous) read pairs for which at least one of the two partners aligns with one of the previously defined genomic intervals is automatically duplicated in the set of other defined intervals in the alignment procedure **(Supplementary Figure S2).**

Since the ambiguity injection procedure is performed under controlled and localized conditions, the quality of the reconstruction proposed by Hicberg is determined by comparing the bins (rows or columns of the contact matrix) relative to twin genomic regions between the initial matrix and the reconstruction resulting from Hicberg. The assessment of the reconstruction is then performed via the computation of Pearson’s correlation of all the genomic bins affected by the injection of ambiguity between the initial vectors and the reconstruction given by Hicberg. This makes it possible to evaluate the closeness of the contact distribution between the initial signal and reconstruction. The closer this correlation coefficient is to 1, the better the reconstruction is. To make this reconstruction score interpretable, we need to compare it with the score obtained via a null model, for which the pairs of twin genomic regions are randomly assigned. Each box plot corresponds to 20 different experiments.

For the performance benchmark, the standard dataset used is a 47 million-read asynchronous *Saccharomyces cerevisiae* Hi-C library generated with a standard protocol (45). To compare the use of RAM memory and CPU computation time, this dataset was subsampled to 1%, 10%, 25%, 50%, 75% and 100%. Both Hicberg and mHiC were tested in their respective standard modes, with the contact matrix resolution set at 2 kb. All programs were run with 16 CPU threads on an Intel(R) Xeon(TM) W-2255 at 3.70 GHz and with 32 Gb of RAM available. The software versions used for this benchmark are Hicberg v0.1.0 and the latest version of mHiC available at https://github.com/yezhengSTAT/mHiC as of January 2025.

## Generation and visualisation of reconstructed contact maps

Chromosomal contact maps without and with reconstruction were normalised via the cooler package and visualised via python code 2whole_matrices_reconstruction_mappability.py.

Theoretical mappability along the chromosomes was computed via the software Genmap (54) https://github.com/cpockrandt/genmap with default parameters, and a bed file was generated for the plot with the contact map.

For the 4C-like signal for the telomeric region of the chimeric chromosome (**Supplementary Figure S25**), the sum of the 5 last bins corresponding to the telomeric region was computed and then plotted along the chromosome (used code: mycoT_4Clike.py).

Correlation matrices were calculated via the correlation_matrice.py code. Briefly, the data were normalized via the balance_cooler function in the cooler package. A detrending relative to the genomic distance was subsequently performed via the distance_law_from_mat function in the hicstuff package. Finally, the correlation matrix is computed via the corrcoef_sparse function in the hicstuff package.

## Tys as 2µ plasmid contact hotspots

For the reconstruction of 2µ plasmid contact signal, a standard Hi-C library in log-phase condition was exploited, and the default parameters of Hicberg were used, as was the standard reference genome (sacCer3). The contact signal of the 2µ-plasmid was computed as previously described (55). Briefly, the 2µ plasmid contact signal is the number of contacts of the 2µ plasmid with a given bin (2 kb resolution) normalized by the number of Hi-C reads of the bin along the genome (to reduce the differences in detectability of certain regions (20)) and divided by the proportion of reads coming from the 2µ sequence (to take into account differences in the number of copies of the plasmid for a given biological condition). The positions of the various transposons were extracted from the SGD database (http://sgd-archive.yeastgenome.org/sequence/S288C_reference/other_features/). Mean plots and contact heatmaps for each element of the different families were generated via homemade python codes <u>plasmid_HSC_1D_agglo.py</u>.

## Tys and cohesin positioning

Reconstruction of Hi-C maps and ChIP-seq of Scc1 protein data were performed with the default parameters of the Hicberg and W303 genomes from the T2T project (W303.asm01.HP0). Annotations of genes and Ty retrotransposons were performed via files from the gff3 annotation file (W303.asm01.HP0.mitochondrial_genome.tidy.gff3 from the https://www.evomicslab.org/db/ScRAPdb database).

The occupancy signals for Scc1 were reconstructed with the Hicberg default parameters of Hicberg. Bedtools was used to convert pair files into bed files, and bedGraphToBigWig from the Short Read Archive suite was used to convert bed files to bigwig files.

The occupancy signal representations for Scc1 for each element of the transposon families and their mean profile were produced via the deeptools tool suite (https://github.com/deeptools/deepTools).

## Identification of the position of transposons from contact signals

A 4C-like plot for a group of genomic elements was generated via custom-made python codes available on GitHub (code: 4Clike_cool_multi.py), e.g., for the group of transposon elements, at each position of the annotated Ty, the 4C-like signal was computed. The calculation of the 4C-like signal is similar to that of the 2µ plasmid contact signal; then, the total signal is summed. If other Ty elements are present, secondary peak(s) can be visualised.

## Experimental confirmation of Ty presence

To validate the presence of an unannotated Ty in the reference genome at genomic position chr12 818,200--818,800 bp, we designated primers on either side of the potential integration (primers used: PL-45 CACTGAAGTTAGTGCCCTTTATGG and PL-46 GGACGTCGGCATAATTGATGGAAC) via the DreamTaq PCR protocol. Sanger sequencing was also performed on the PCR products to confirm the presence of the transposon.

## Declarations

### Availability of data and materials

The software and documentation are available at https://github.com/koszullab/Hicberg. This implementation works with commonly paired-end data in fastq format. All the data associated with this study are publicly available, and their reference numbers are listed in Supplementary Table 1. All the scripts used to reproduce the figures and analyses are available at https://github.com/axelcournac/Hicberg_bioanalysis. The Hicberg algorithm (1.0.1) has also been deposited on Zenodo <u>DOI: 10.5281/zenodo.15594031</u>.

## Acknowledgments

All the members of the Spatial Regulation of Genomes unit are thanked for stimulating discussions and feedback, especially Amaury Bignaud, Fabien Girard and Jacques Sérizay. This work used the computational and storage services (TARS cluster) provided by the IT department at Institut Pasteur, Paris. We would like to thank Stéphane Descorps Declere, Johann Dreo, Pascale Lesage, Nicolas Maillet and Julien Mozziconacci for their stimulating discussions and Agnès Thierry for advice on the experimental section.

## Contributions

SG and AC conceptualized the method. SG implemented the method in Python. SG and AC applied this method to the biological data. SG and AC interpreted the biological results. SO and MD helped with the code implementation. PL performed the experimental validation. SG and AC wrote the manuscript with feedback from all the authors. Supervision and funding acquisition: AC.

## Corresponding authors

Please send correspondence to Axel Cournac.

## Ethics declarations

**Ethics approval and consent to participate**

Not applicable.

## Consent for publication

Not applicable.

## Supplementary information

Additional file 1: Reconstruction of genomic signals from repeated elements.

