## Supplementary data for "Hicberg: Reconstruction of contact signals from repeated elements"

### Supplementary Figures

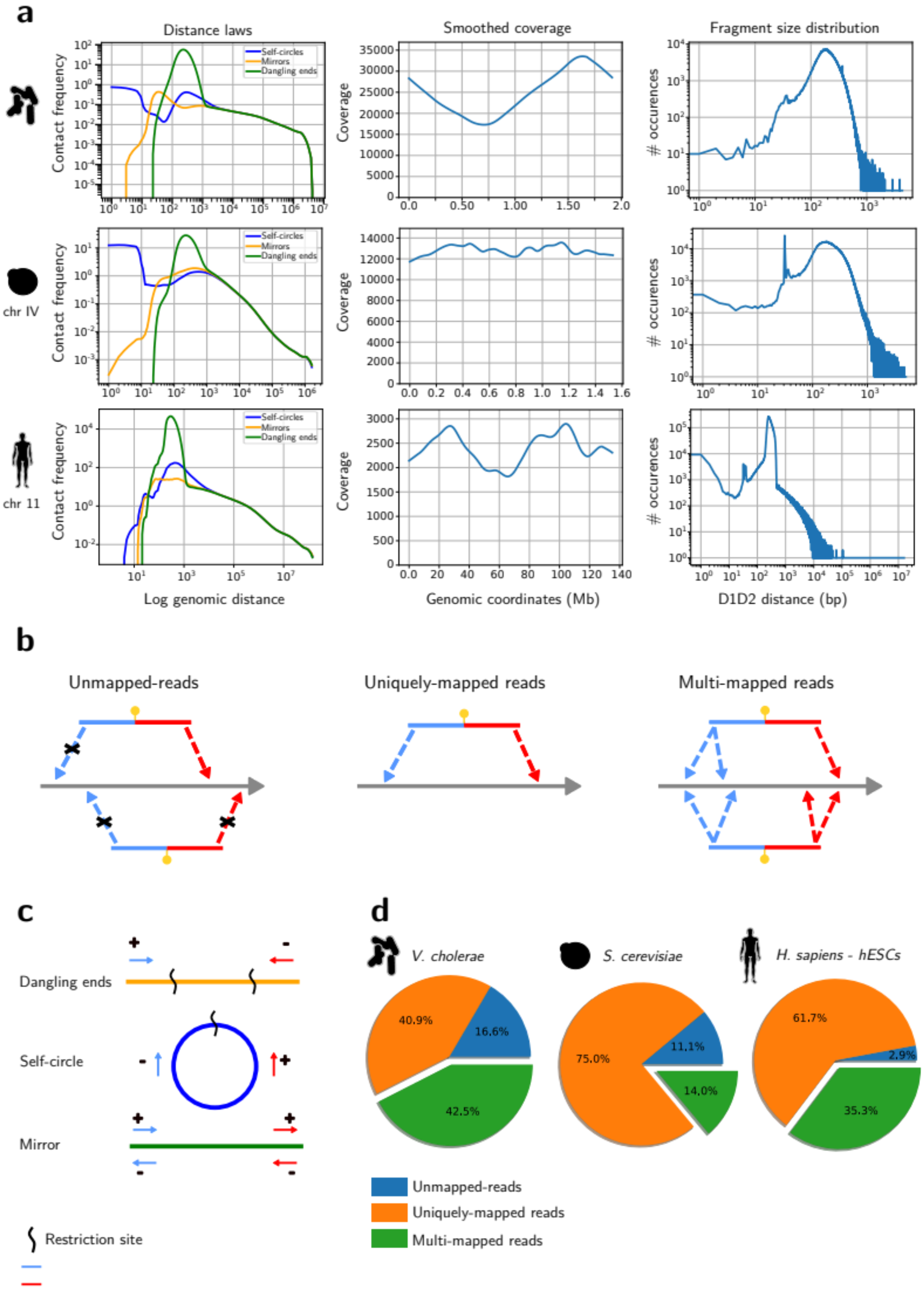


**Figure S1. Leveraging Hi-C library biases for accurate assignment of multi-mapping reads.**

**a** Distributions of Hi-C event types (self-circles, mirrors, and dangling ends) over genomic distance, genomic coverage along the genome, and D1D2 fragment size distributions for

*Vibrio cholerae*, *Saccharomyces cerevisiae* chromosome IV, and human chromosome 11. he distinct profiles for each species highlight the potential for using these features to discriminate the origin of multi-mapping reads. **b** Schematic representation of possible Hi-C read mapping configurations: unmapped (at least one read mate unmapped), uniquely mapped (both mates map uniquely), and multi-mapped (at least one mate maps to multiple locations). **c** Illustration of Hi-C events: dangling ends (incomplete restriction enzyme digestion), self-circles (DNA folding back on itself; orientation: -+), and mirrors (incorrect end orientations; orientation: ++ or --). These events, characterized by mate orientations and undigested fragments, contribute to the distinctive profiles observed in (a). **d** Pie charts showing the distribution of mapping configurations (unmapped, uniquely mapped, multi-mapped) for *Vibrio cholerae*, *Saccharomyces cerevisiae*, and human (hESCs) genomes. The proportion of multi-mapping reads varies across species and genome complexity, emphasizing the need for accurate assignment to fully capture chromosome architecture.

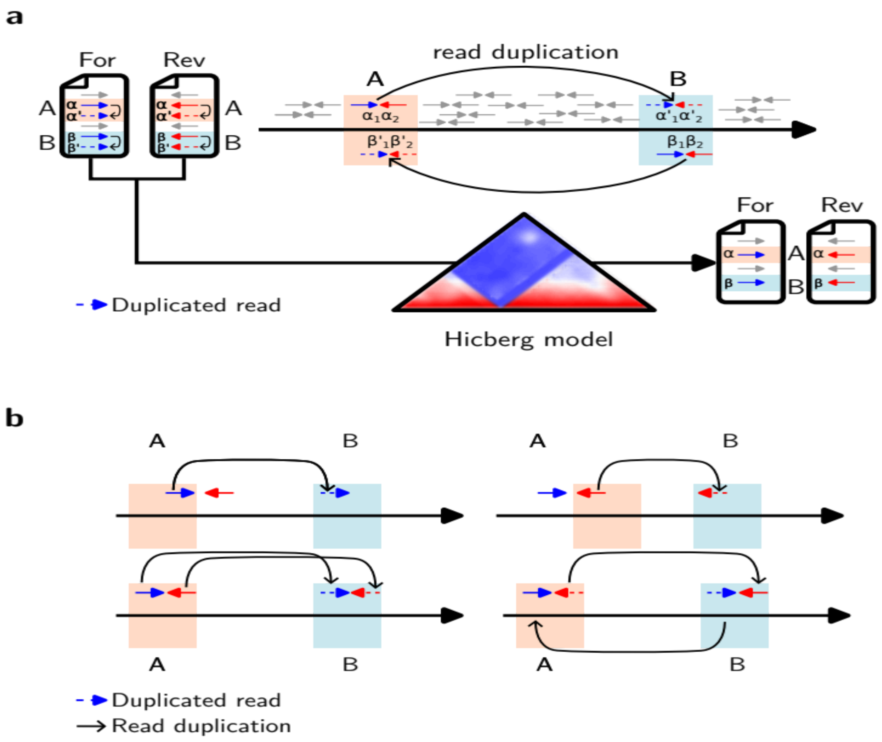


**Figure S2. Evaluation of Hicberg's performance using simulated repeated elements.**

**a** Schematic overview of the simulation strategy. Repeated elements are simulated by duplicating reads mapping to selected genomic intervals (A and B). Hicberg then assigns these artificially multi-mapping reads based on the statistical profiles of uniquely mapped reads. The accuracy of Hicberg's assignments is evaluated by comparing the reconstructed contact map to the ground truth (uniquely mapped reads before duplication).

**b** Detailed illustration of the read duplication process. Reads with at least one mate uniquely mapping to interval A or B are selected. The alignment of the mate within A or B is then duplicated in the other interval, creating artificial multi-mapping reads. For example, a read pair with the forward mate in A and the reverse mate outside A and B will have the forward mate duplicated in B, resulting in two possible pairs. This process simulates the ambiguity introduced by repeated elements in real Hi-C data.

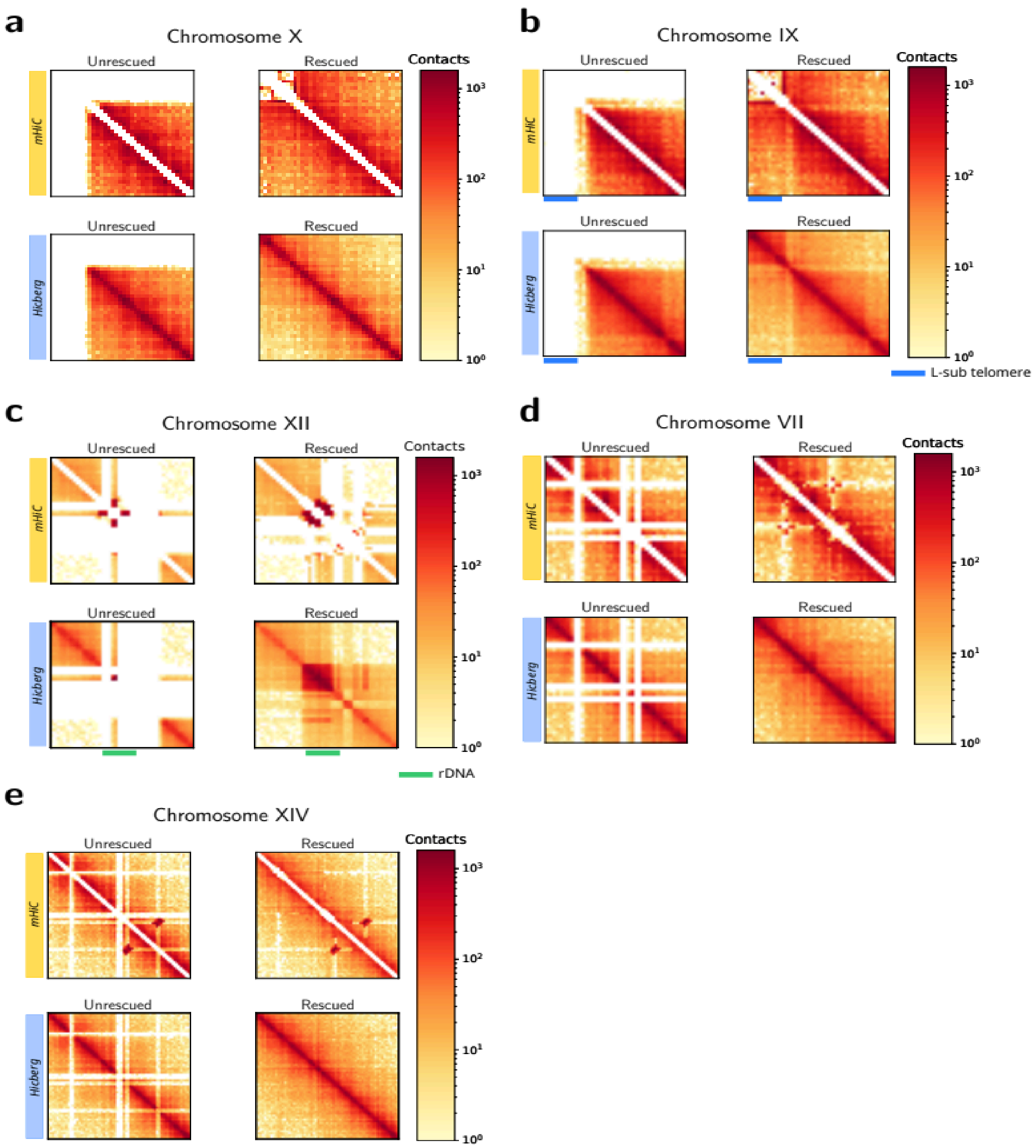


**Figure S3. Comparison of reconstructions from Hicberg and mHiC in *Saccharomyces cerevisiae*.**

This figure compares the performance of Hicberg and mHiC in reconstructing Hi-C contact maps for regions of the *Saccharomyces cerevisiae* genome containing repetitive elements. **a** Subtelomeric region of chromosome X. Hicberg (bottom) generates a complete and coherent contact map, while mHiC (top) produces a hollow diagonal and misses contacts in the subtelomeric region. **b** Subtelomeric region of chromosome IX. Similar to (a),

Hicberg (bottom) provides a more comprehensive reconstruction compared to mHiC (top). **c** rDNA tandem repeat region on chromosome XII. Hicberg (bottom) accurately captures the structure of the rDNA region, whereas mHiC (top) shows a hollow diagonal and an empty region corresponding to the rDNA. **d,** Region with repeated sequences on chromosome VII. Hicberg (bottom) reconstructs a coherent contact map, while mHiC (top) exhibits a hollow diagonal and spurious artifacts at mid-range distances. **e,** Region with repeated sequences on chromosome XIV. Again, Hicberg (bottom) provides a more accurate and complete reconstruction compared to mHiC (top), which shows a hollow diagonal and artifactual interactions.

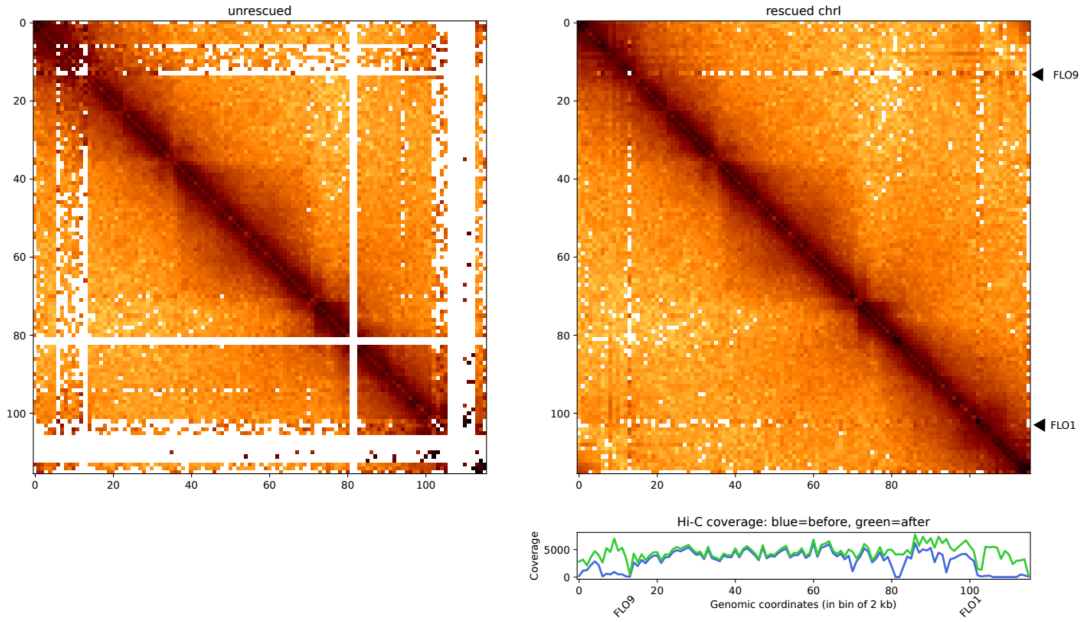


**Figure S4: Contact map of Chromosome I of *Saccharomyces cerevisiae*** without and with Hicberg reconstruction with Hi-C coverages below right (bin = 2 kb, with normalisation). Under-covered bins after reconstruction correspond to the *FLO9* and *FLO1* genes.

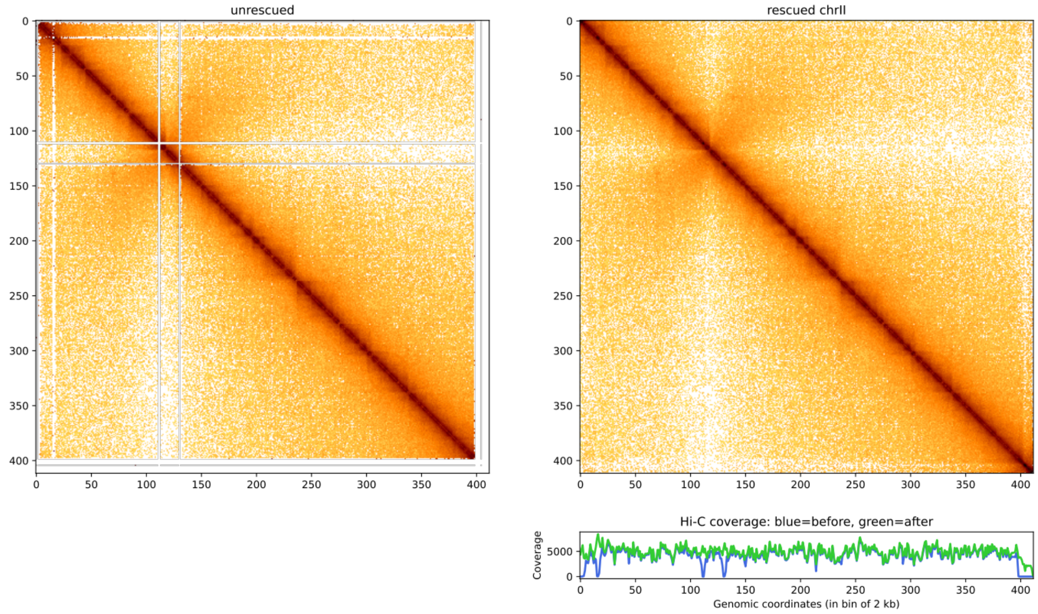


**Figure S5: Contact map of Chromosome II of *Saccharomyces cerevisiae*** without and with Hicberg reconstruction with Hi-C coverages below right (bin = 2 kb, with normalisation).

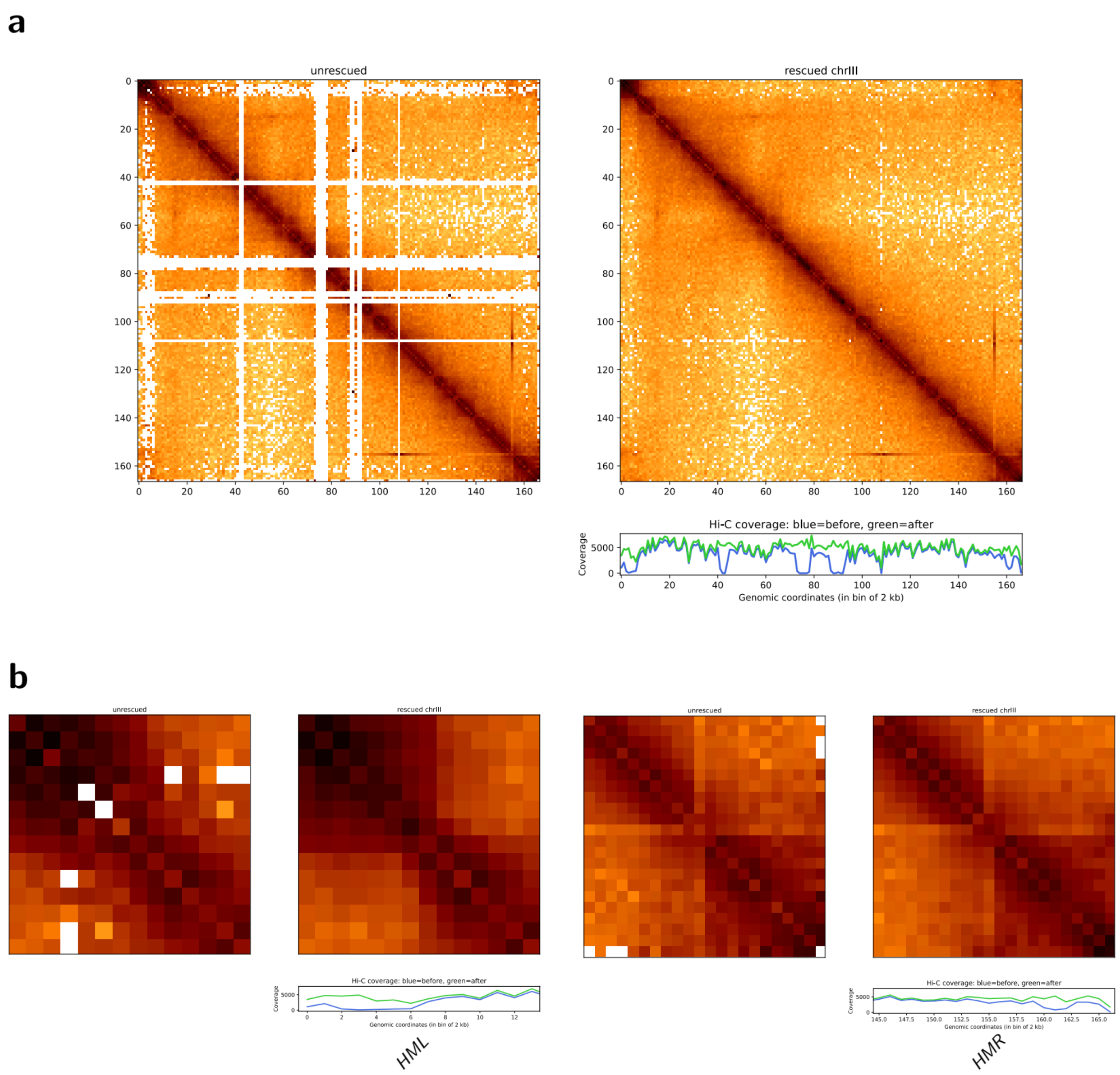


**Figure S6: a Contact map of Chromosome III of *Saccharomyces cerevisiae*** without and with Hicberg reconstruction with Hi-C coverages below right (bin = 2 kb, with normalisation).

**b** Zoom on HML and HMR loci showing borders patterns.

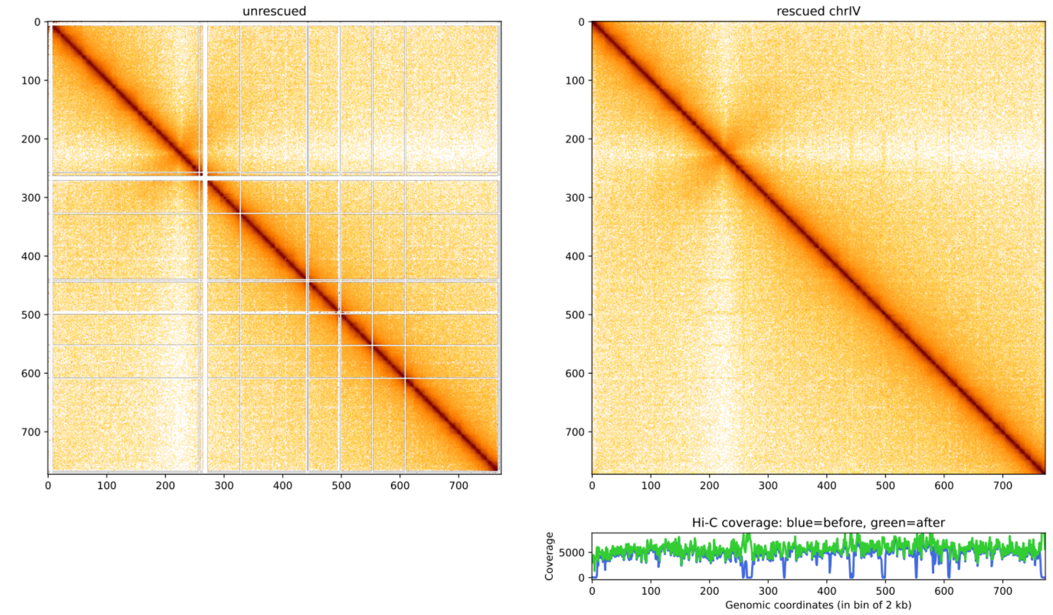


**Figure S7: Contact map of Chromosome IV of *Saccharomyces cerevisiae*** without and with Hicberg reconstruction with Hi-C coverages below right (bin = 2 kb, with normalisation).

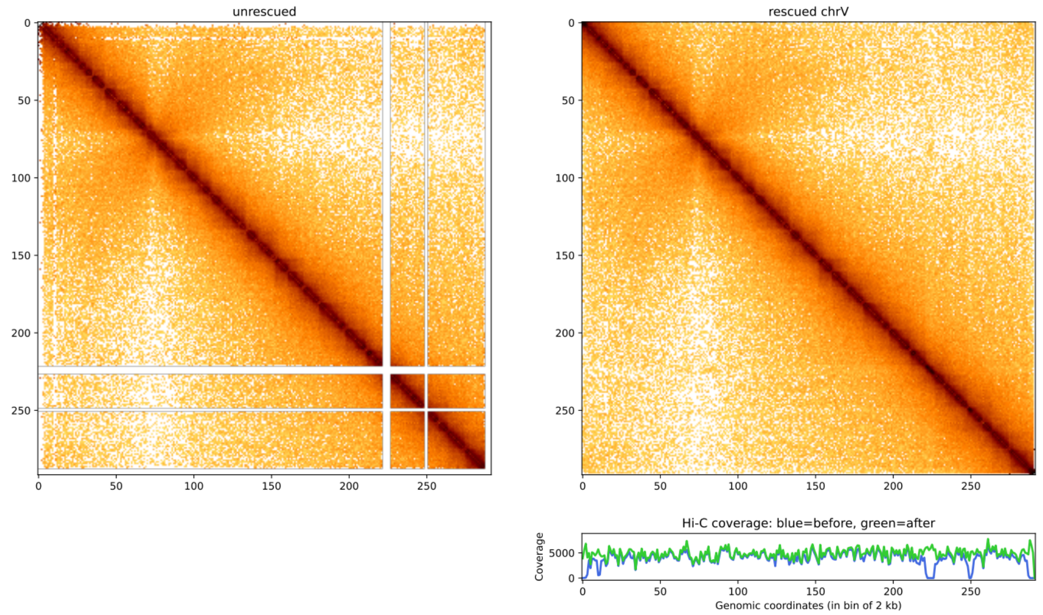


**Figure S8: Contact map of Chromosome V of *Saccharomyces cerevisiae*** without and with Hicberg reconstruction with Hi-C coverages below right (bin = 2 kb, with normalisation).

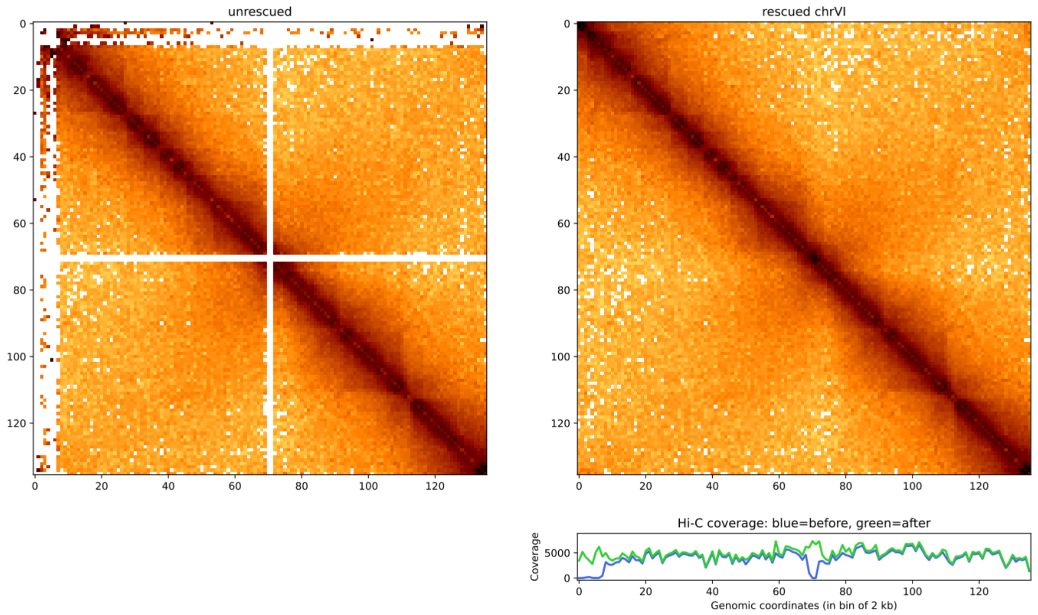


**Figure S9: Contact map of Chromosome VI of *Saccharomyces cerevisiae*** without and with Hicberg reconstruction with Hi-C coverages below right (bin = 2 kb, with normalisation).

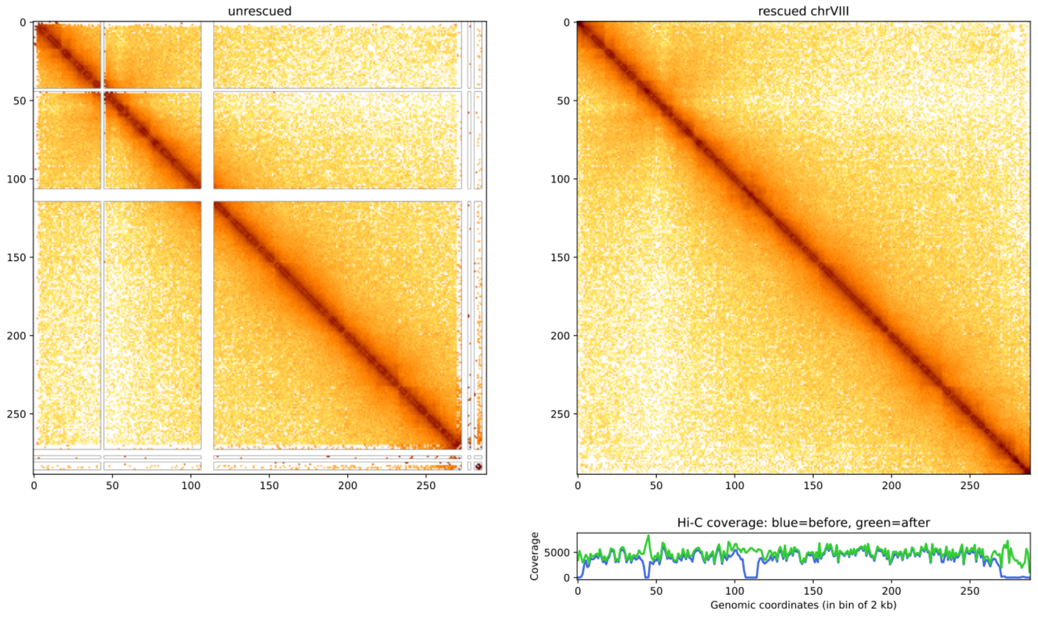


**Figure S10: Contact map of Chromosome VIII of *Saccharomyces cerevisiae*** without and with Hicberg reconstruction withHi-C coverages below right (bin = 2 kb, with normalisation).

**a**

**b**

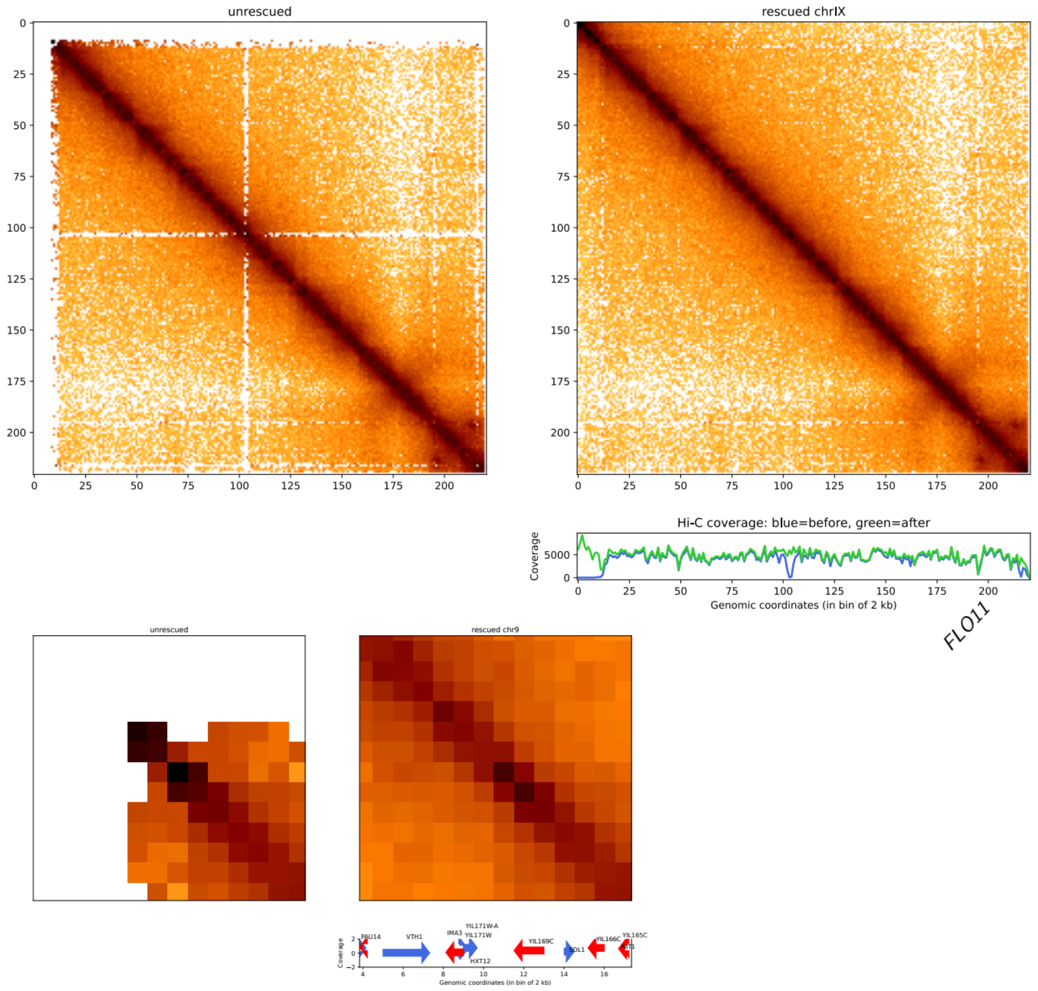


**Figure S11: A, Contact map of Chromosome IX of *Saccharomyces cerevisiae*** without and with Hicberg reconstruction with Hi-C coverages below right.

**b** Zoom on TEL9L region showing border pattern between HXT12 and YIL169C (CSS1) genes.

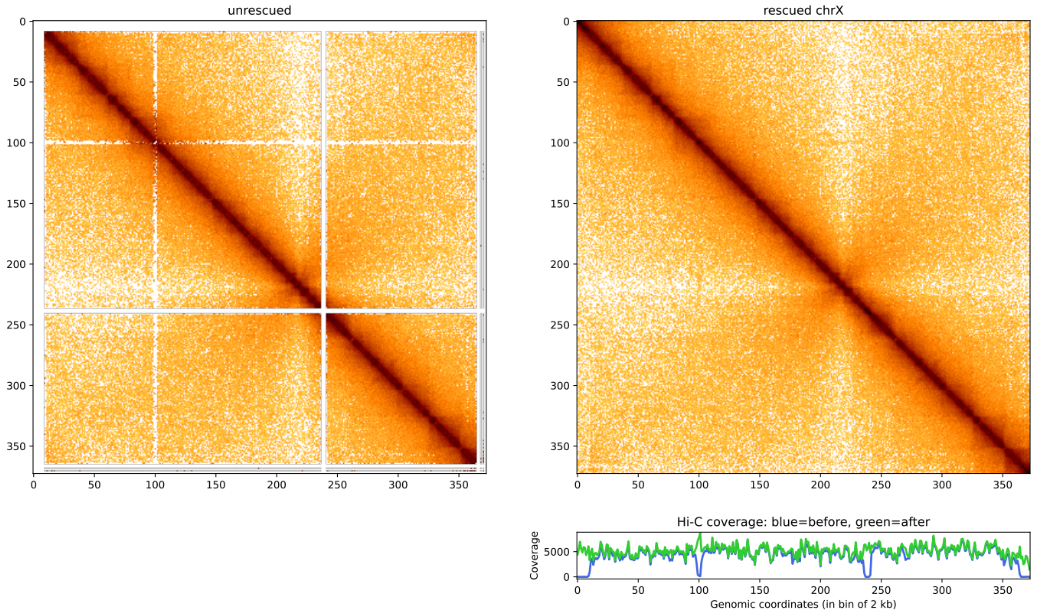


**Figure S12: Contact map of Chromosome X of *Saccharomyces cerevisiae*** without and with Hicberg reconstruction with Hi-C coverages below right (bin = 2 kb, with normalisation).

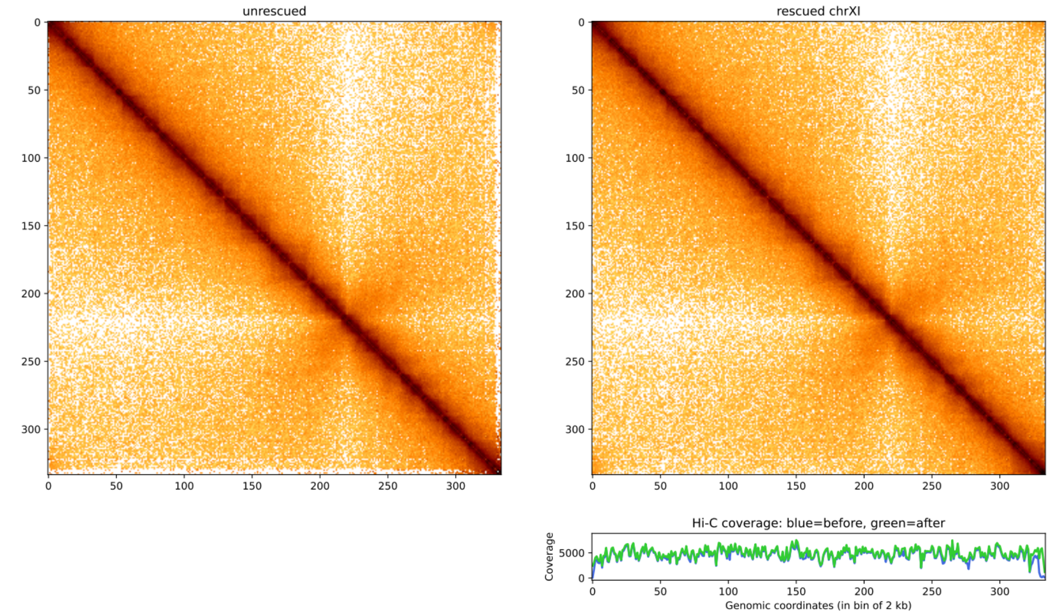


**Figure S13: Contact map of Chromosome XI of *Saccharomyces cerevisiae*** without and with Hicberg reconstruction with Hi-C coverages below right (bin = 2 kb, with normalisation).

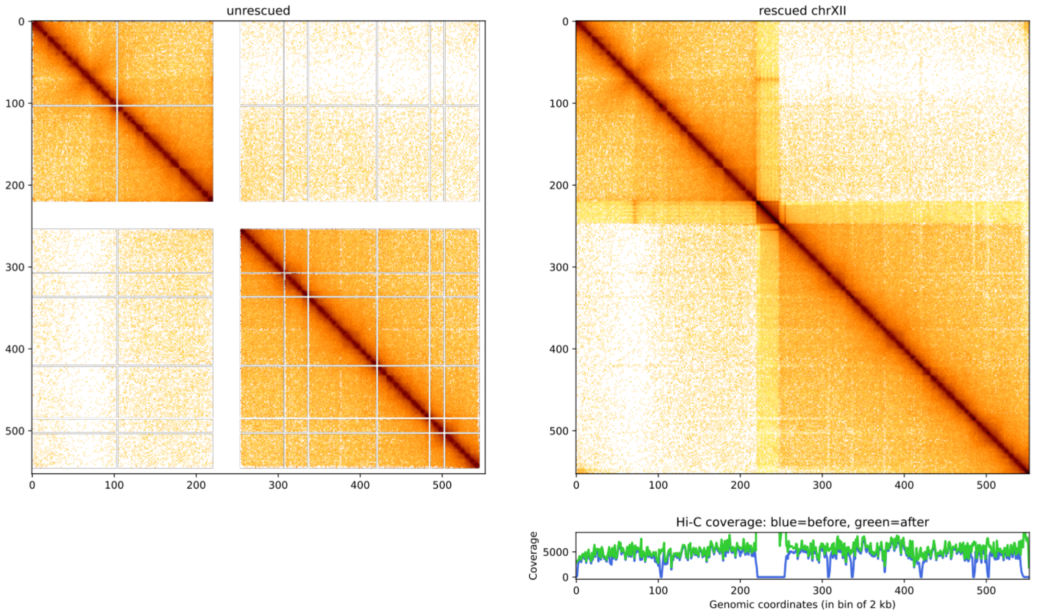


**Figure S14: Contact map of Chromosome XII of *Saccharomyces cerevisiae*** without and with Hicberg reconstruction with Hi-C coverages below right (bin = 2 kb, with normalisation).

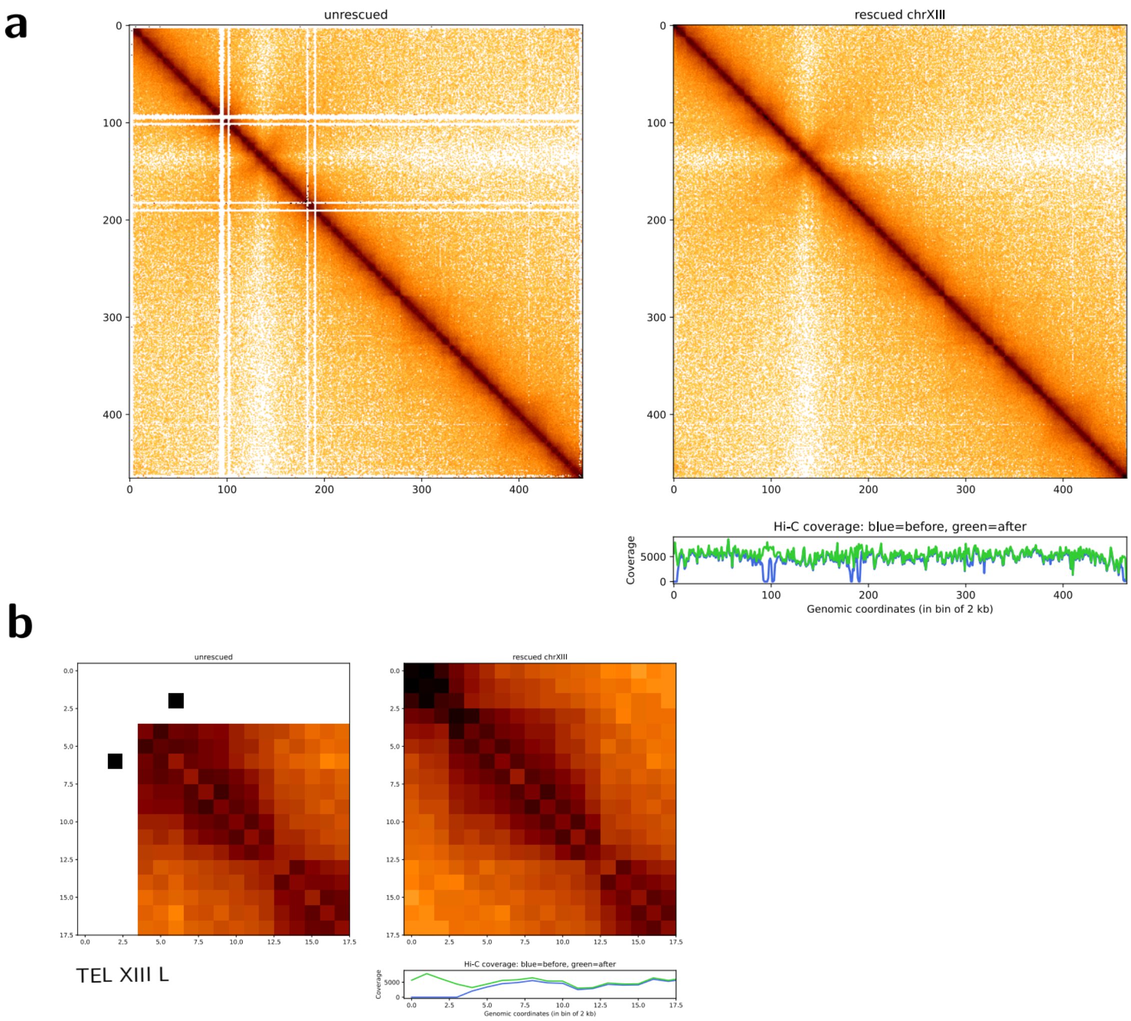


**Figure S15: a, Contact map of Chromosome XIII of *Saccharomyces cerevisiae*** without and with Hicberg reconstruction with Hi-C coverages below right (bin = 2 kb, with normalisation).

**b,** Zoom on TEL XIII Left region showing a border pattern after Hicberg reconstruction.

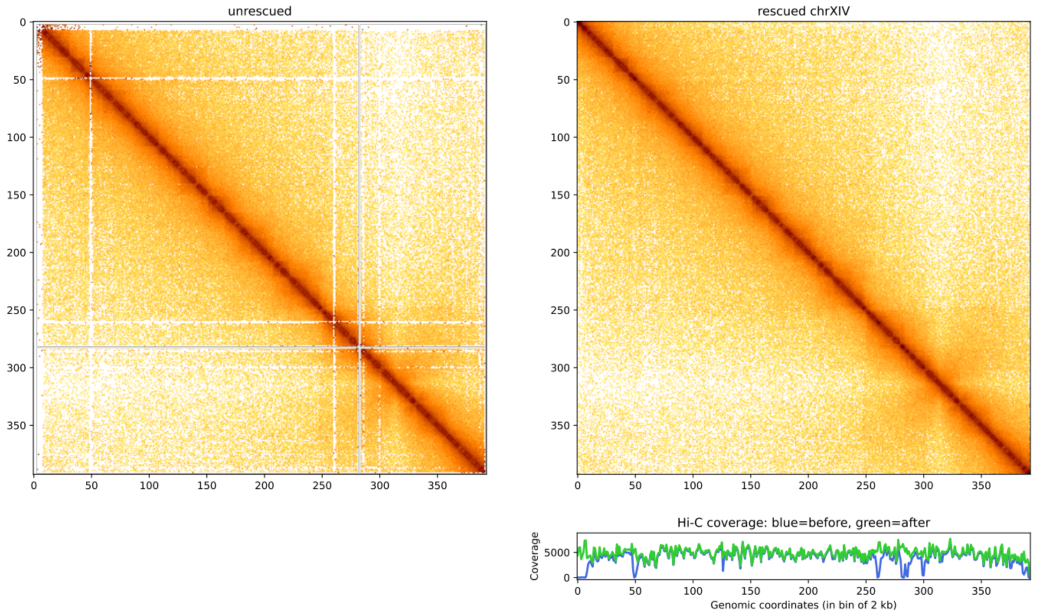


**Figure S16: Contact map of Chromosome XIV of *Saccharomyces cerevisiae*** without and with Hicberg reconstruction with Hi-C coverages below right (bin = 2 kb, with normalisation).

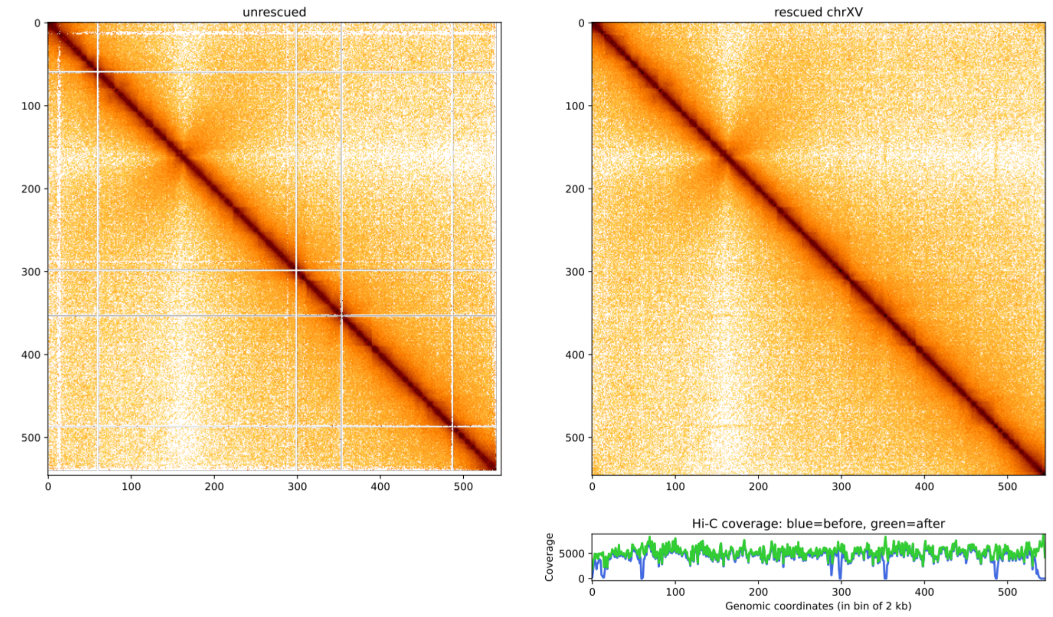


**Figure S17: Contact map of Chromosome XV of *Saccharomyces cerevisiae*** without and with Hicberg reconstruction with Hi-C coverages below right (bin = 2 kb, with normalisation).

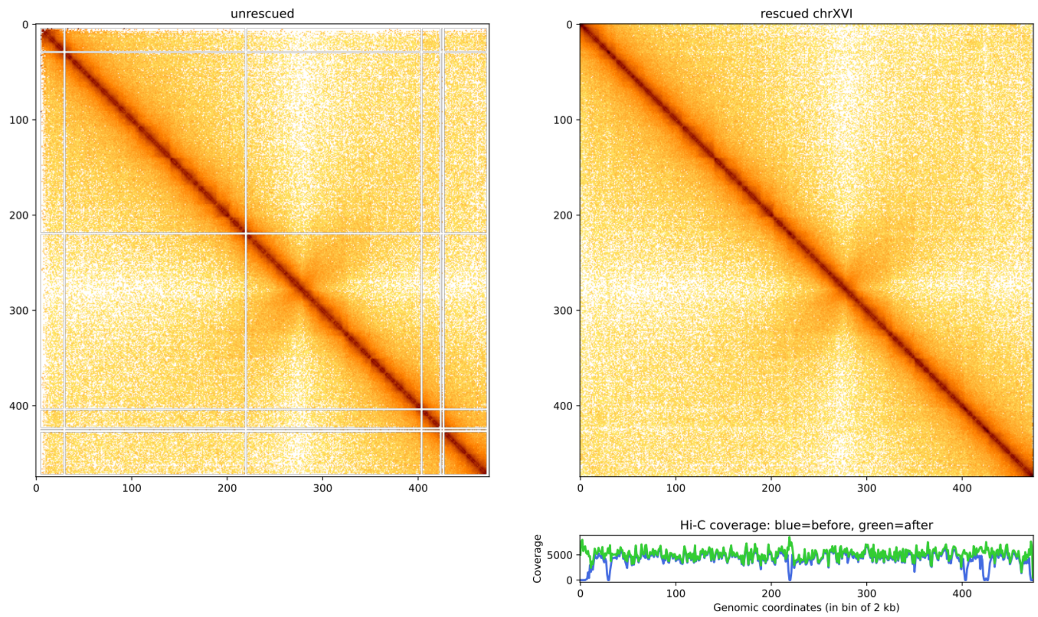


**Figure S18: Contact map of Chromosome XVI of *Saccharomyces cerevisiae*** without and with Hicberg reconstruction with Hi-C coverages below right (bin = 2 kb, with normalisation).

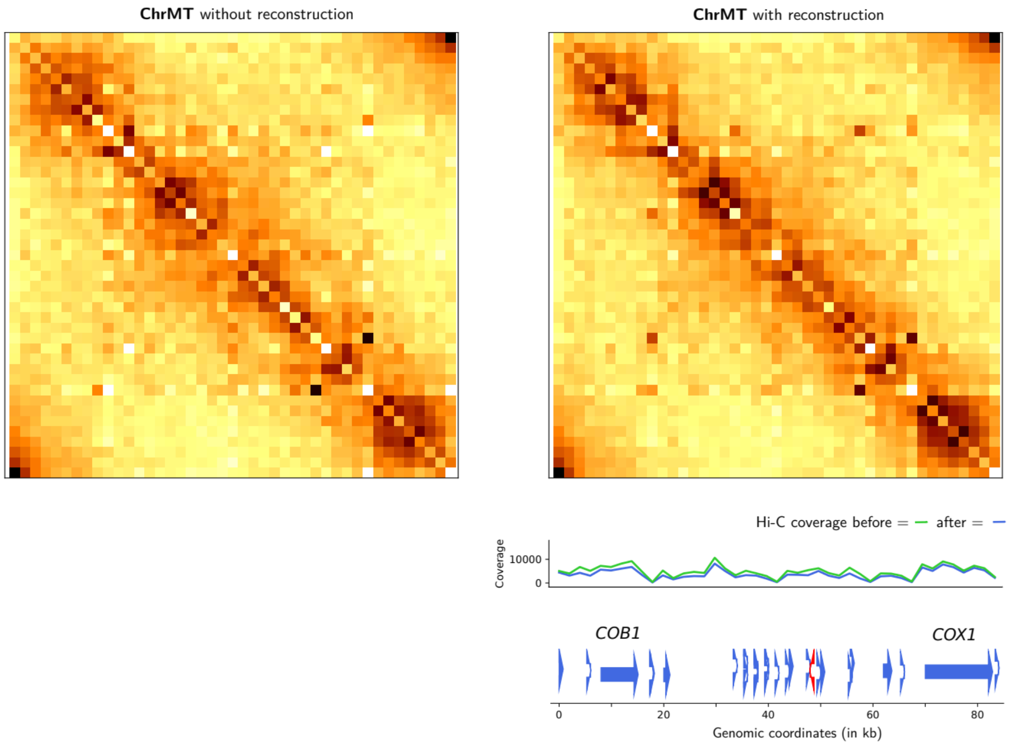


**Figure S19: Contact map of Mitochondrion of *Saccharomyces cerevisiae*** without and with Hicberg reconstruction with Hi-C coverages et gene annotation below right (bin = 2 kb, with normalisation).

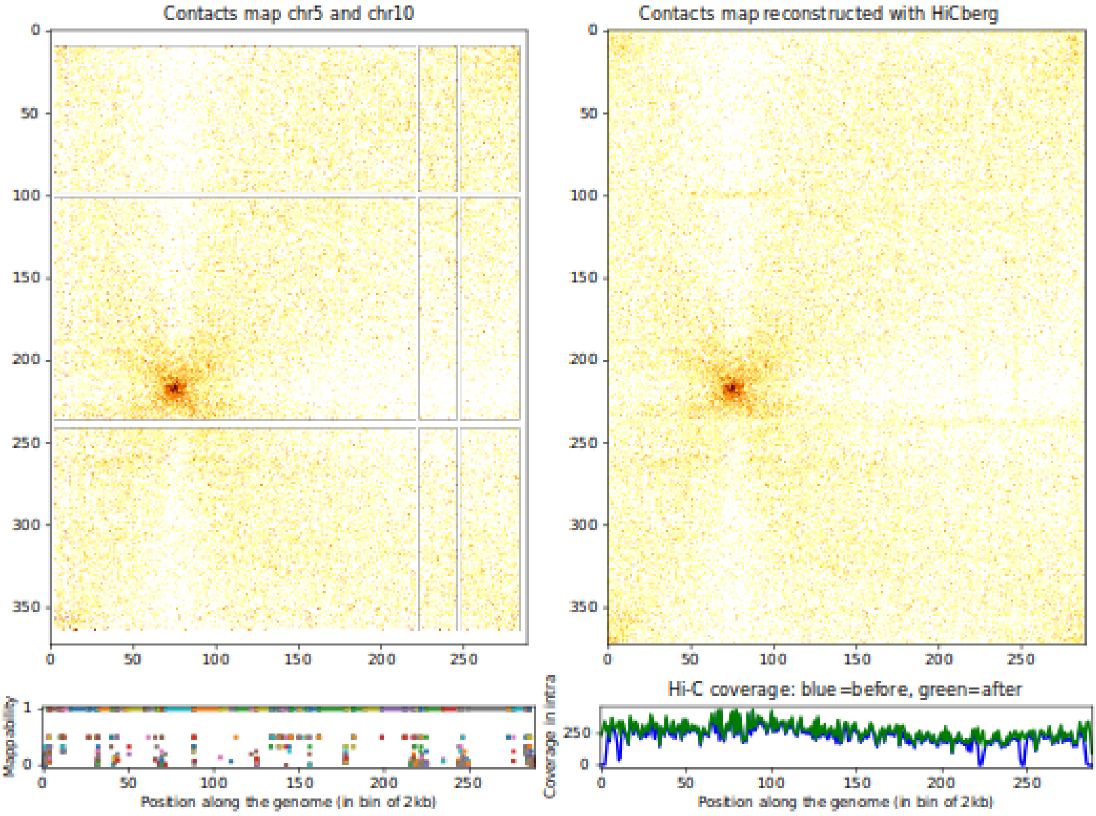
Contacts map chr5 and chr10
Contacts map reconstructed with HiCberg
Mappability
Position along the genome (in bin of 2kb)
Hi-C coverage: blue=before, green=after
Coverage in intra

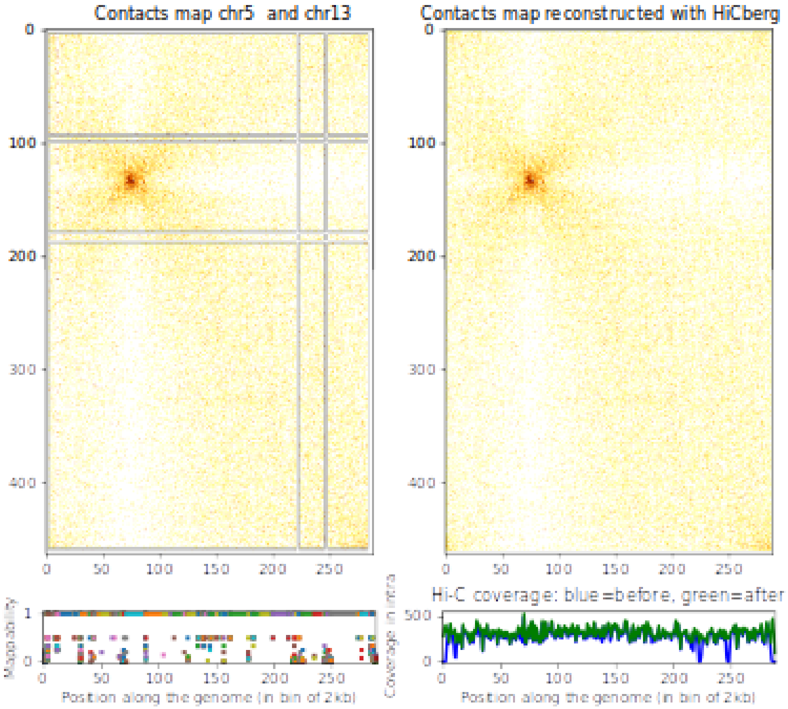
Contacts map chr5 and chr13
Contacts map reconstructed with HiCberg
Mappability
Position along the genome (in bin of 2kb)
Hi-C coverage: blue=before, green=after
Coverage in intra

**Figure S20: Examples of contact maps between different chromosomes of *Saccharomyces cerevisiae*** without and with Hicberg reconstruction with mappabililty signal below left and Hi-C coverages below right (bin = 2 kb, with normalisation).

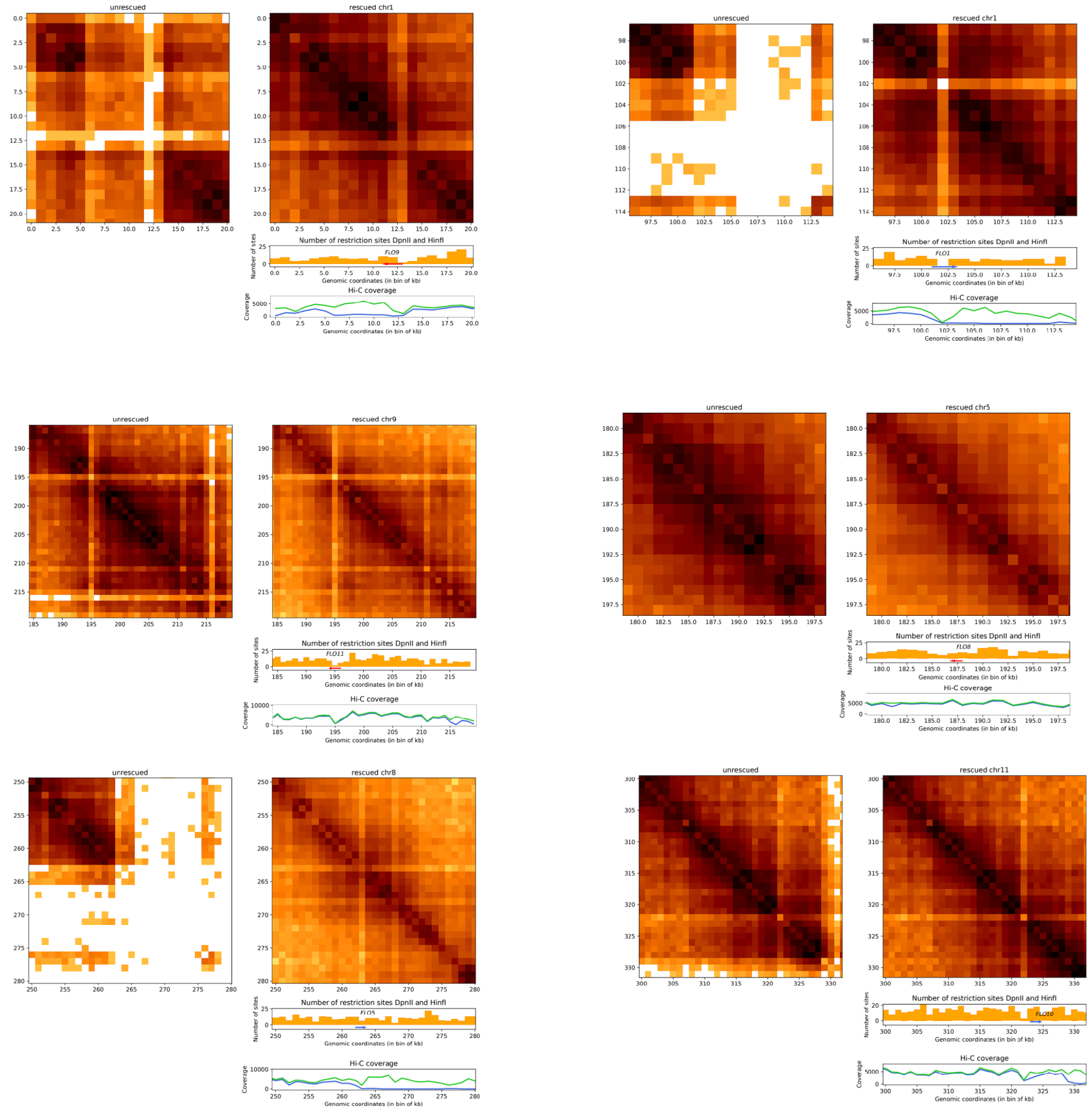


**Figure S21: Raw contact maps of the 6 FLO genes in *Saccharomyces cerevisiae*** without and with Hicberg reconstruction with histograms of the number of restriction sites (DpnII, HinfI) used in the Hi-Cprotocol and coverages below right (bin = 2 kb, raw data). All the genes showed a deficit in Hi-C coverage even after reconstruction mainly due to the low number of restriction sites (DpnII, HinfI) used in the Hi-C protocol. Only the *FLO8* gene on chr5 showed normal coverage.

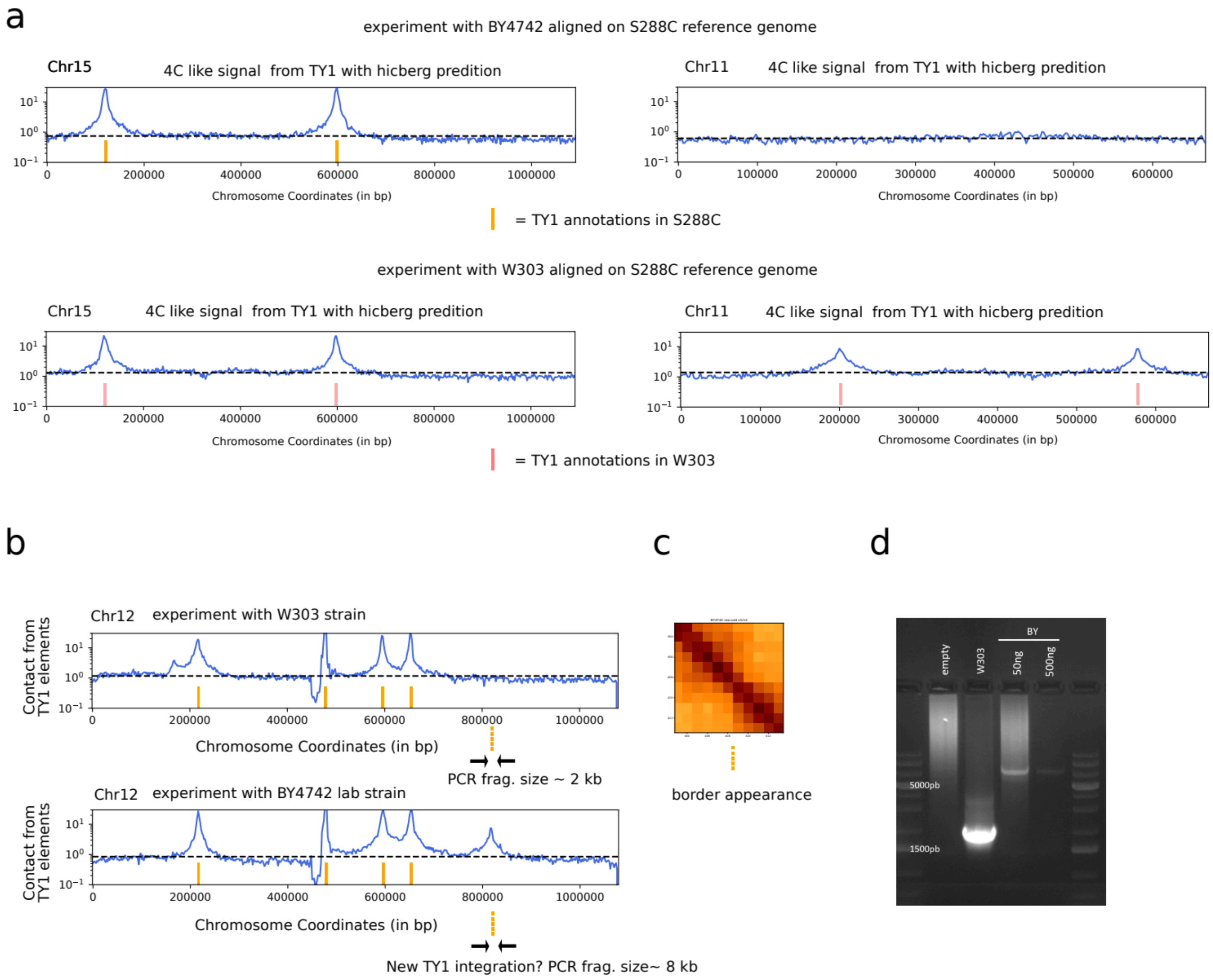


**Figure S22. Identification of a TY1 element in *Saccharomyces cerevisiae* S288C using Hicberg compared to the reference genome GCF_000146045.2, sacCer3.**

**a** Virtual 4C profiles of aggregated TY1 elements on chromosomes XV and XI in S288C (top) and W303 (bottom) strains. Peaks in the virtual 4C profiles correspond to known TY1 annotations. The absence of a peak on chromosome XI in S288C correlates with the lack of annotated TY1 elements in this region. **b** Virtual 4C profiles of aggregated TY1 elements on chromosome XII in W303 (top) and S288C (bottom) strains. While peaks in W303 align with known TY1 annotations, S288C exhibits an additional peak with a similar profile (orange, dotted), suggesting a TY1 insertion. **c** Zoom in Hi-C contact map of the suspected insertion site on chromosome XII in S288C. A boundary pattern is visible, co-localizing with the predicted TY1 insertion site identified in (b)**. d** PCR validation of the

TY1 insertion. Electrophoresis gel showing PCR products amplified from W303 and S288C using primers flanking the predicted insertion site. A band of approximately 8kb, consistent with the size of TY1, is present in S288C (third and fourth lanes) but absent in W303 (second lane), confirming the presence of the TY1 insertion which was confirmed by Sanger sequencing (data not shown).

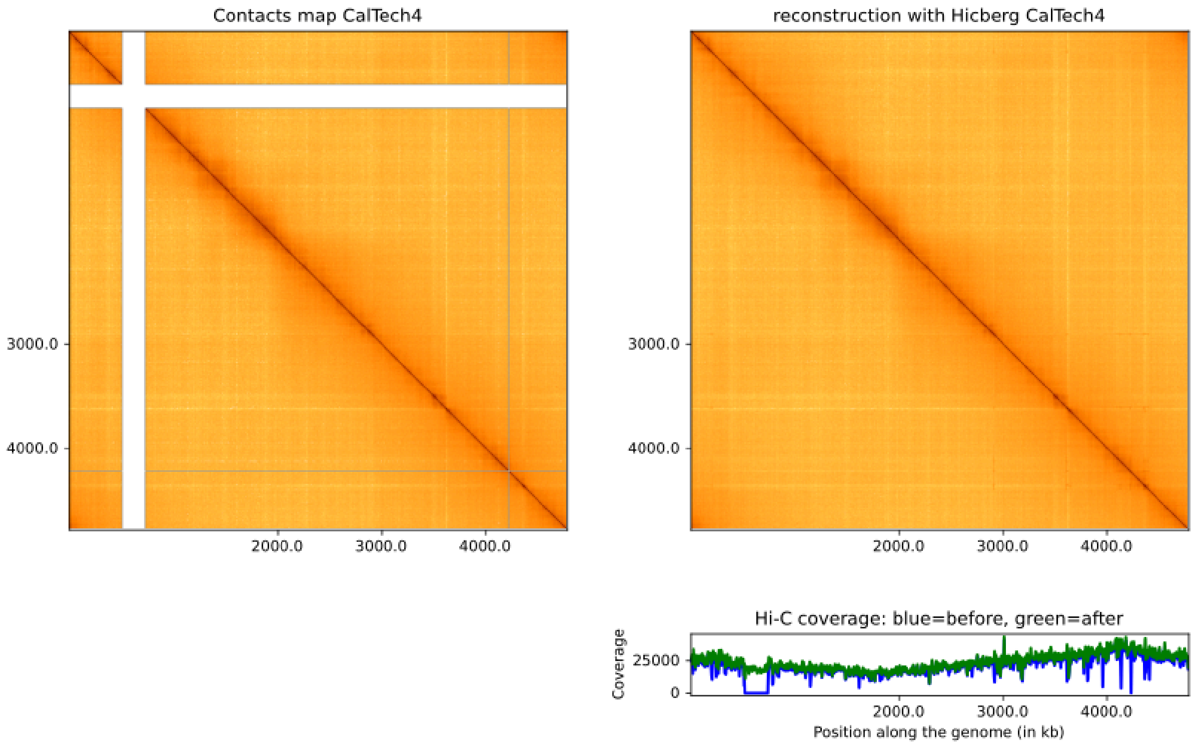


**Figure S23: Contact maps of *Escherichia coli* containing a duplicated region** without and with Hicberg reconstruction with Hi-C coverages below right, bin = 5 kb.

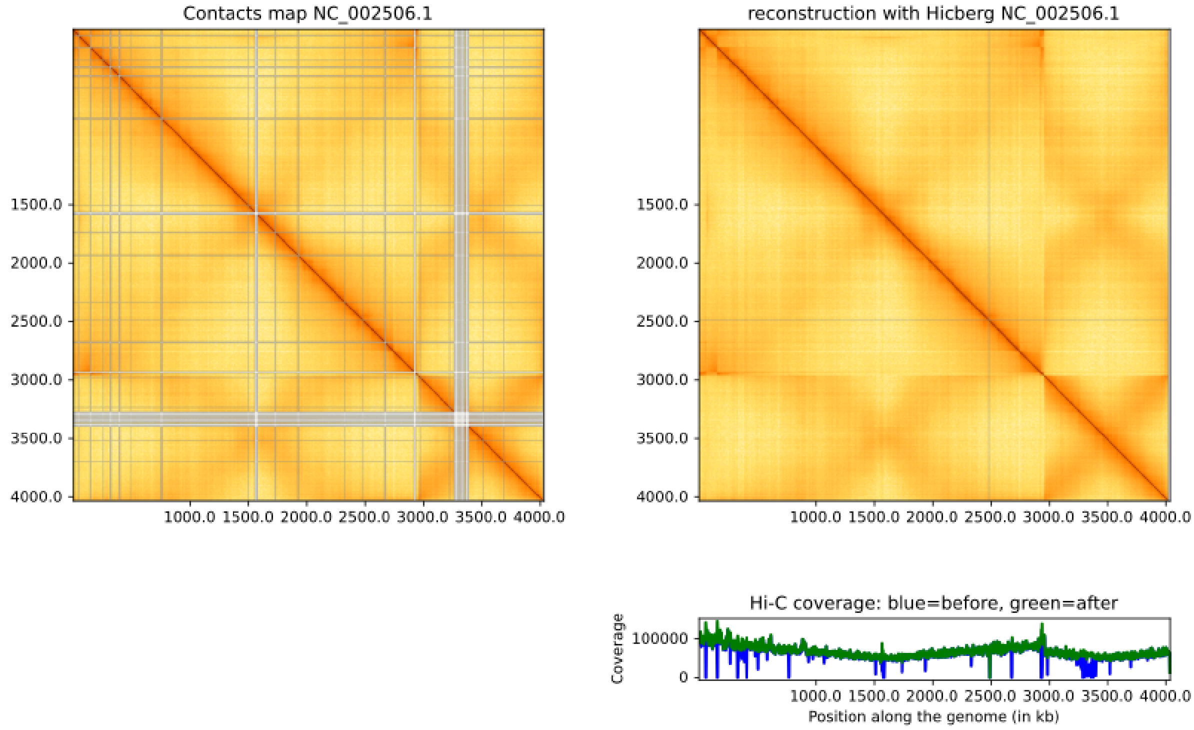


**Figure S24: Contact maps of** ***Vibrio Cholerae*** without and with Hicberg reconstruction with Hi-C coverages below right, bin = 2 kb.

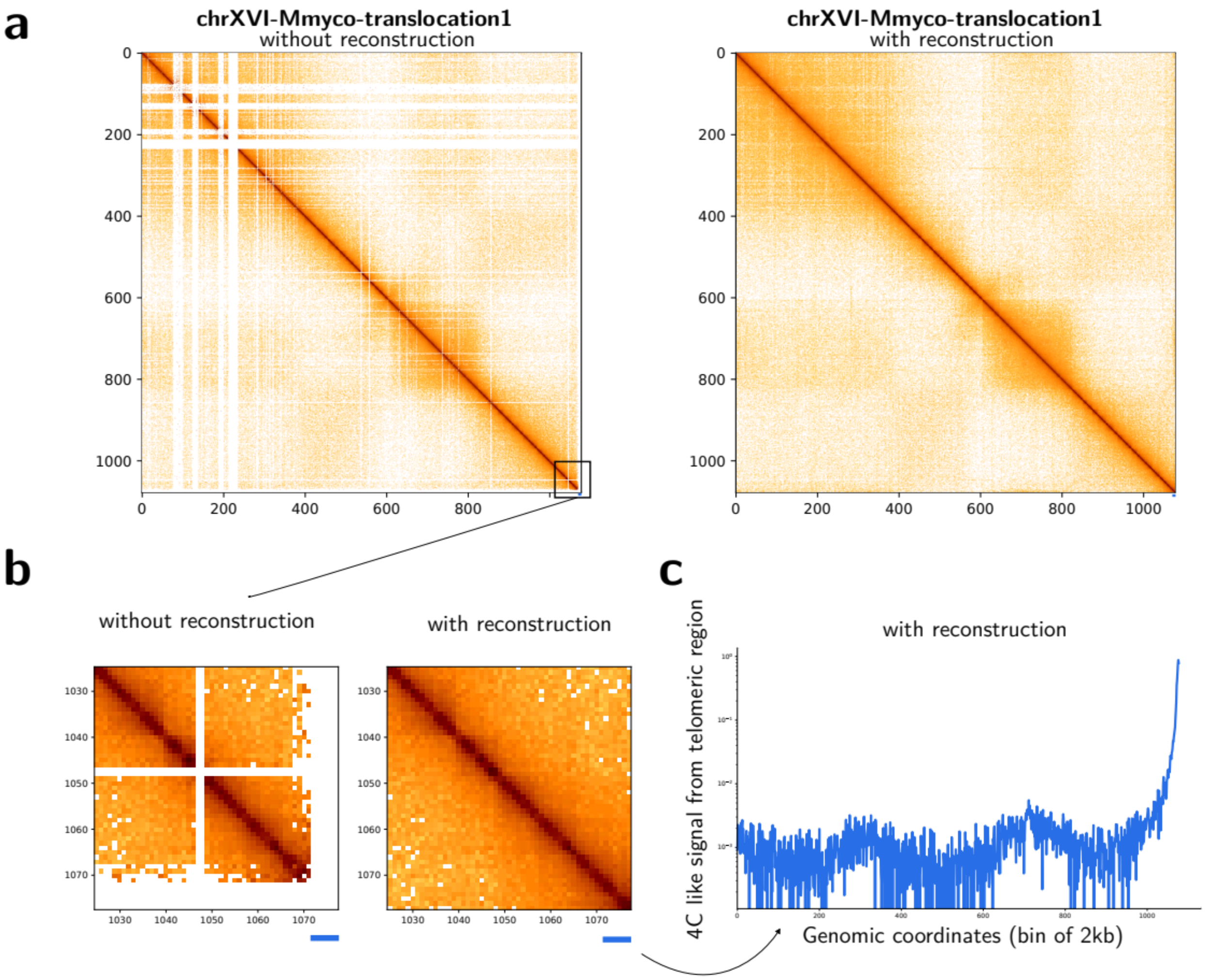


**Figure S25: Contact maps of chimeric chromosome *S. cerevisiae* and M. *mycoides***
**a** Contact map of chimeric chrXVI-Mmyco translocation 1 chromosome without and with Hicberg reconstruction (bin = 2 kb, with normalisation). **b** Zoom on the telomeric region without and with Hicberg reconstruction (bin = 2 kb, with normalisation). **c** 4C like signal from the telomeric region (identified by a blue rectangle in a and b) only present in the reconstructed data (bin = 2 kb, with normalisation) showing the organisation into compartments.

**Figure S26.**

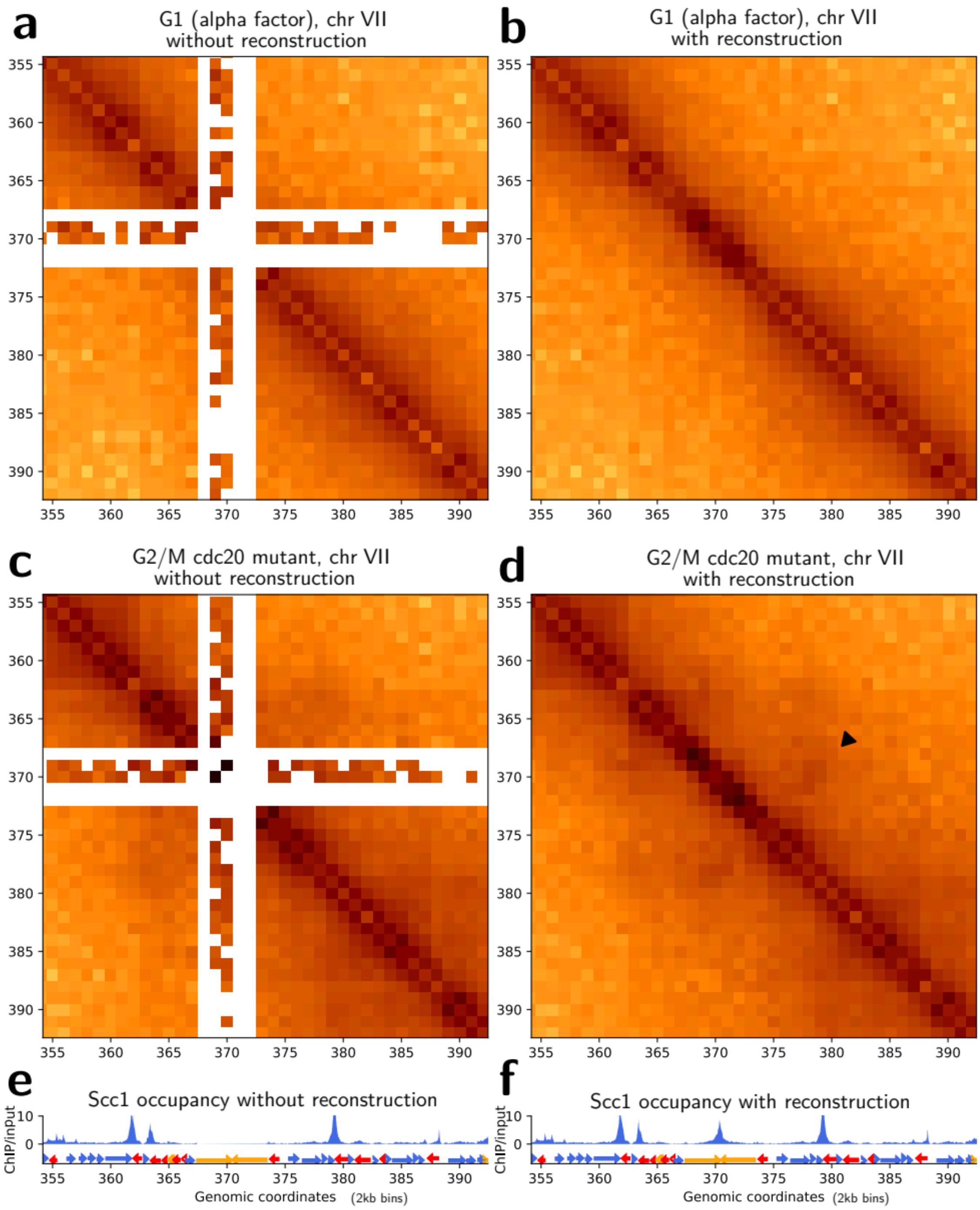


**Positionning of cohesin and TY retrotransposons in *S. cerevisiae* genome**

**a, b** Zooms of contact map of chromosome VII of *S. cerevisiae* without and with reconstruction, bin = 2 kb in a population synchronized in G1 phase (W303 strain, alpha factor, data from Piazza et al., 2021). **c,d** Zoom of contact map of chromosome VII of *S. cerevisiae* without and with reconstruction, bin = 2 kb in a population synchronized in G2/M phase (W303 strain, cdc20 mutant, data from Piazza et al., 2021). **e, f** Cohesin occupancy signal from ChIP-seq of Scc1 in G2/M phase without and with reconstruction. Forward and reverse genes are represented by blue and red arrows respectively. Yeast transposons (TY) are represented by orange arrows.

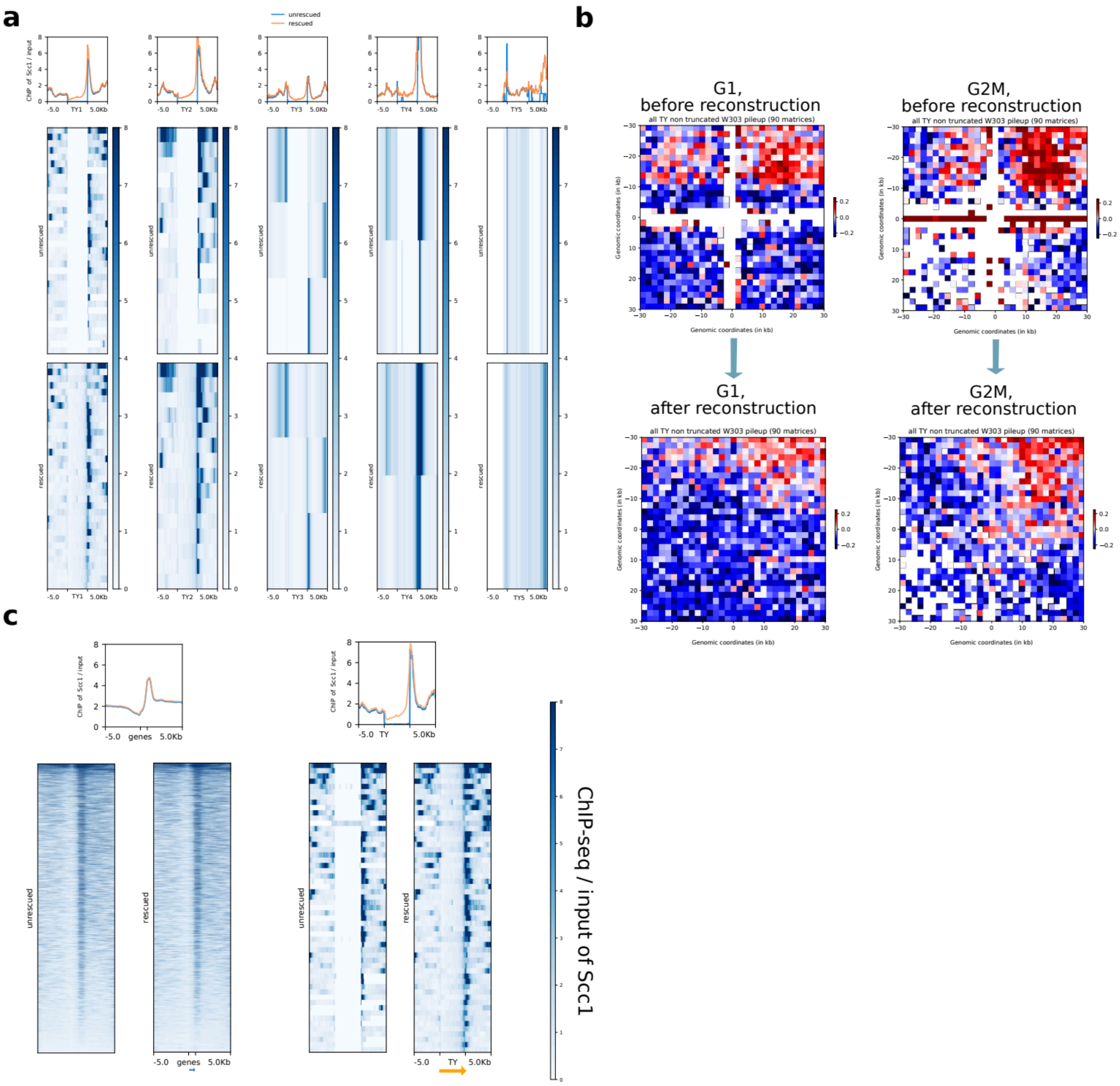


**Figure S27. Averaged behavior of yeast transposons in ChIP-seq of Scc1 and Hi-C**

**a** Agglomerated plot of cohesin occupancy signal (ChIP-seq of Scc1 in G2/M phase) for the 5 retrotransposons families (TY1, TY2, TY3, TY4, TY5) without and with Hicberg reconstruction.

**b** Agglomerated plot between different TY elements from same chromosomes in G1 and G2/M phases without and with Hicberg reconstruction.

**c** Agglomerated plot of cohesin occupancy signal (ChIP-seq of Scc1 in G2/M phase) for all genes and all yeast transposons elements (TY) without and with Hicberg reconstruction in the W303 genetic background, biological replicate.

| Experiment type and condition | Specie | Figure | Ref | Identifier |
|---|---|---|---|---|
| Hi-C, WT of *S. cerevisiae* S288C (BY4741), asynchronous population, grown at 25 °C | *Saccharomyces cerevisiae* | **Fig 3, Fig 5**<br>**Fig. S3 to S22** | This work | AC1 |
| HiC, WT of *E.coli* using MluCI and HpaII | *Escherichia coli* | **Fig S23** | (Cockram *et al.*, 2021) | SRR10394901 |
| Hi-C, WT of *V. cholerae O1*, log phase using HpaII | *Vibrio cholerae* | **Fig S24** | (Cockram *et al.*, 2021) | SRR10394900 |
| Hi-C, XVIfMmycot1 strain +DMSO, rep2 (G1) | *Saccharomyces cerevisiae* | **Fig 4** | (Meneu *et al*, 2025) | SRR31398213 |
| Hi-C, WT of *E. festucae,* hyphae were grown in axenic culture | *Epichloe festucae* | **Fig 4** | (Winter *et al.*, 2018) | SRR8238190 |
| Hi-C, WT of *S. cerevisiae* W303, G1 arrest | *Saccharomyces cerevisiae* | **Fig 6** | (Piazza *et al*, 2021) | SRR12284705 |
| Hi-C, WT of *S. cerevisiae* W303, G2/M arrest with the mutant cdc20, replicate 1 | *Saccharomyces cerevisiae* | **Fig 6** | (Piazza *et al*, 2021) | SRR13736508 |
| ChIP-seq of cohesin, W303 Scc1-PK9 DSB t4 + nocodazole (IP) | *Saccharomyces cerevisiae* | **Fig 6** | (Piazza *et al*, 2021) | SRR15041199 |
| ChIP-seq of cohesin, W303 Scc1-PK9 DSB t4 + nocodazole (Input) | *Saccharomyces cerevisiae* | **Fig 6** | (Piazza *et al*, 2021) | SRR15041197 |
| ChIP-seq of Scc1-HA FB218-4a Noco IP | *Saccharomyces cerevisiae* | **Fig 7** | (Chapard et al. 2025) | CH376 |
| ChIP-seq of Scc1-HA FB218-4a Noco Input | *Saccharomyces cerevisiae* | **Fig 7** | (Chapard et al. 2025) | CH381 |
| ChIP-seq of Scc1-HA FB218-4a Noco IP | *Saccharomyces cerevisiae* | **Fig S27** | (Chapard et al. 2025) | CH377 |
| ChIP-seq of Scc1-HA FB218-4a Noco Input | *Saccharomyces cerevisiae* | **Fig S27** | (Chapard et al. 2025) | CH382 |

**Supplementary Table S1: Datasets generated or reanalyzed in the present study.**
The last column indicates either the identifier for the raw reads available on the Short Read Archive server (SRA) (https://www.ncbi.nlm.nih.gov/sra) or on Gene Expression Omnibus server (GEO) (https://www.ncbi.nlm.nih.gov/geo).
